# Glutamate inputs send prediction error of reward but not negative value of aversive stimuli to dopamine neurons

**DOI:** 10.1101/2023.11.09.566472

**Authors:** Ryunosuke Amo, Naoshige Uchida, Mitsuko Watabe-Uchida

## Abstract

Midbrain dopamine neurons are thought to signal reward prediction errors (RPEs) but the mechanisms underlying RPE computation, particularly contributions of different neurotransmitters, remain poorly understood. Here we used a genetically-encoded glutamate sensor to examine the pattern of glutamate inputs to dopamine neurons. We found that glutamate inputs exhibit virtually all of the characteristics of RPE, rather than conveying a specific component of RPE computation such as reward or expectation. Notably, while glutamate inputs were transiently inhibited by reward omission, they were excited by aversive stimuli. Opioid analgesics altered dopamine negative responses to aversive stimuli toward more positive responses, while excitatory responses of glutamate inputs remained unchanged. Our findings uncover previously unknown synaptic mechanisms underlying RPE computations; dopamine responses are shaped by both synergistic and competitive interactions between glutamatergic and GABAergic inputs to dopamine neurons depending on valences, with competitive interactions playing a role in responses to aversive stimuli.

## Introduction

A fundamental computation that the brain performs is to compare its expectations and reality. It has been proposed that the difference between expectations and reality (called prediction errors) is the driving force behind perception and learning^1–5^. Although prediction errors can be defined with simple mathematical formulas, how intricate networks of neurons compute prediction errors remains largely elusive.

The activity pattern of dopamine neurons in the lateral ventral tegmental area (VTA) is relatively uniform^6,7^, and dopamine responses have long been quantitatively formalized in reinforcement learning theory as reward prediction error (RPE), the discrepancy between actual and expected reward^4,8^. According to this theory, RPE can be used to learn to take actions in a specific “state”, a distinct combination of environmental and internal information, to maximize future rewards. Because of the simplicity of the learning rule—learning from surprising outcomes, which is consistent with animal behaviors^9,10^—RPE is widely used to model reinforcement learning in diverse situations^11^.

While the RPE characteristics of dopamine neuron activity have been intensively studied, the mechanism of how RPE is computed based on inputs to dopamine neurons is not well understood. Building on the simple mathematical formula of RPE, i.e. subtraction of expected reward value from actual reward value, many models assume that a specific brain area sends information about actual reward while a different brain area sends information about expected reward to dopamine neurons^4,12–16^. VTA dopamine neurons receive input from multiple types of neurons, including from glutamate and GABA neurons^17–21^. Given the excitatory and inhibitory nature of these forms of neurotransmissions, respectively, many models proposed that dopamine neurons might implement this subtraction by combining glutamate reward signals and GABA expectation signals^12^. However, recordings from monosynaptic inputs to dopamine neurons revealed that information for both reward and expectation is distributed and mixed in single presynaptic neurons across various brain regions^22^. The previous study showed that dopamine neurons uniquely signal relatively more “complete” RPE that is consistent across different task events, whereas activity patterns of presynaptic neurons are more diverse and correspond only to the “partial” RPE^22^. These results suggested that RPE may be partially computed in multiple nodes in the neural circuits, not only at the level of dopamine neurons.

On the other hand, disinhibition of GABA inputs has often been proposed to generate dopamine phasic activation because of powerful and stringent gating roles of disinhibition as seen in the basal ganglia^23^. A series of studies identified the lateral habenula (lHb) and its projection target, the rostromedial tegmental area (RMTg), as an important pathway which conveys computed RPE signals to dopamine neurons^21,24–28^. Neurons in both lHb and RMTg signal RPE in an opposite sign than dopamine neurons, which suggests that the direct projection from GABA neurons in RMTg to dopamine neurons can theoretically produce RPE signals in dopamine neurons^25–28^. However, it was reported that excitation of DA neurons precedes inhibition of lHb neurons^28^, and that ablation of RMTg neurons caused specific deficits in responses to aversive events in dopamine neurons, rather than completely removing RPE signals^26^. Thus, while RMTg GABA inputs send inverse RPE to dopamine neurons, other inputs, likely distributed, play an important role in shaping RPE signals in dopamine neurons.

In this study, we examined the population activity of glutamate inputs to dopamine neurons, focusing on characteristics of temporal difference (TD) errors. Classic studies have proposed that dopamine activity patterns reflect TD errors, a specific form of RPE that has proven especially useful in machine learning^4,29^. TD errors are computed locally between adjacent states at each time step to signal changes in reward expectation. This process supports incremental learning to predict reward value at gradually earlier time points. A recent study confirmed a fundamental prediction of this theory, namely, a gradual temporal shift of dopamine responses to earlier and earlier time points between cue and reward^30^. While information about reward and expectation are distributed in presynaptic neurons, it is not known how glutamate and GABA inputs are used to generate TD errors in dopamine neurons. The common idea, which directly applies the TD error computation in theory to the circuit model is that both glutamate and GABA neurons carry expectation signals with slight time lags. The inhibitory expectation signals are then subtracted from excitatory expectation signals in dopamine neurons to compute derivatives^12,14^, although this does not match the observation that RMTg GABA neurons already signal RPE^25–27^. In the present study, we found that the activity pattern of glutamate inputs to dopamine neurons show characteristics of TD errors but differ in important ways from the signals conveyed by dopamine neurons. Specifically, glutamate inputs were excited by aversive stimuli, while DA neurons in the same areas were inhibited. This activation, coupled with GABA inputs which are reported to excite at aversive stimuli^25,26,31^, determines whether DA neurons are excited or inhibited by an aversive stimulus in situations. Opioids altered this excitation-inhibition balance probably through the μ-opioid receptor-enriched GABA input pathway^21,32–34^. Our results demonstrate both redundancy and division of labor by glutamate and GABA inputs, suggesting a strategy to overcome neural constraints to form bi-directional RPE signals in dopamine neurons.

## Result

### Detection of glutamate release at dopamine neurons

Dopamine neurons receive input, both excitatory and inhibitory, from various areas in the brain^17–19,21,35–38^. To examine information conveyed specifically by excitatory input to dopamine neurons, we used a genetically encoded glutamate sensor. We injected adeno-associated virus (AAV) to express a recently improved glutamate sensor (SFiGluSnFR)^39^ in the ventral tegmental area (VTA) of dopamine transporter (DAT)-Cre mice (Figures 1A and 1B). The glutamate sensor was specifically expressed in dopamine neurons and distributed throughout cell bodies and dendritic fibers (Figure 1B). To monitor the glutamate release at dopamine neurons, an optical fiber was implanted into the VTA for fiber fluorometry. To monitor dopamine release simultaneously, dopamine sensor (GrabDA2m)^40^ was expressed in the ventral striatum (VS) and its fluorescent signals were monitored through an optical fiber implanted in the same location. We targeted an area of VS for recording where dopamine release typically signals reward prediction error (RPE)^30,41^. Overall, simultaneous recording of dopamine release at VS and glutamate release onto dopamine neurons showed similar activity patterns including strong activation at water reward, which was absent in control fluorescence signals (Figure 1C). A significant increase of activity at the water reward delivery was observed in both dopamine release and glutamate input activity (Figure 1D). The water-evoked responses were faster in glutamate sensor signals (rise: 250±17.1 ms, decay: 488±46.4 ms) compared to dopamine sensor signals (rise: 387±41.0 ms, decay: 1222±81.1 ms) (Figure 1E). These results indicate that the natural fluctuation of dopamine sensor signals and glutamate sensor signals are correlated at water reward, and that the temporal resolution of the glutamate sensor signals is sufficient to compare with dopamine sensor signals.

**Figure 1.**
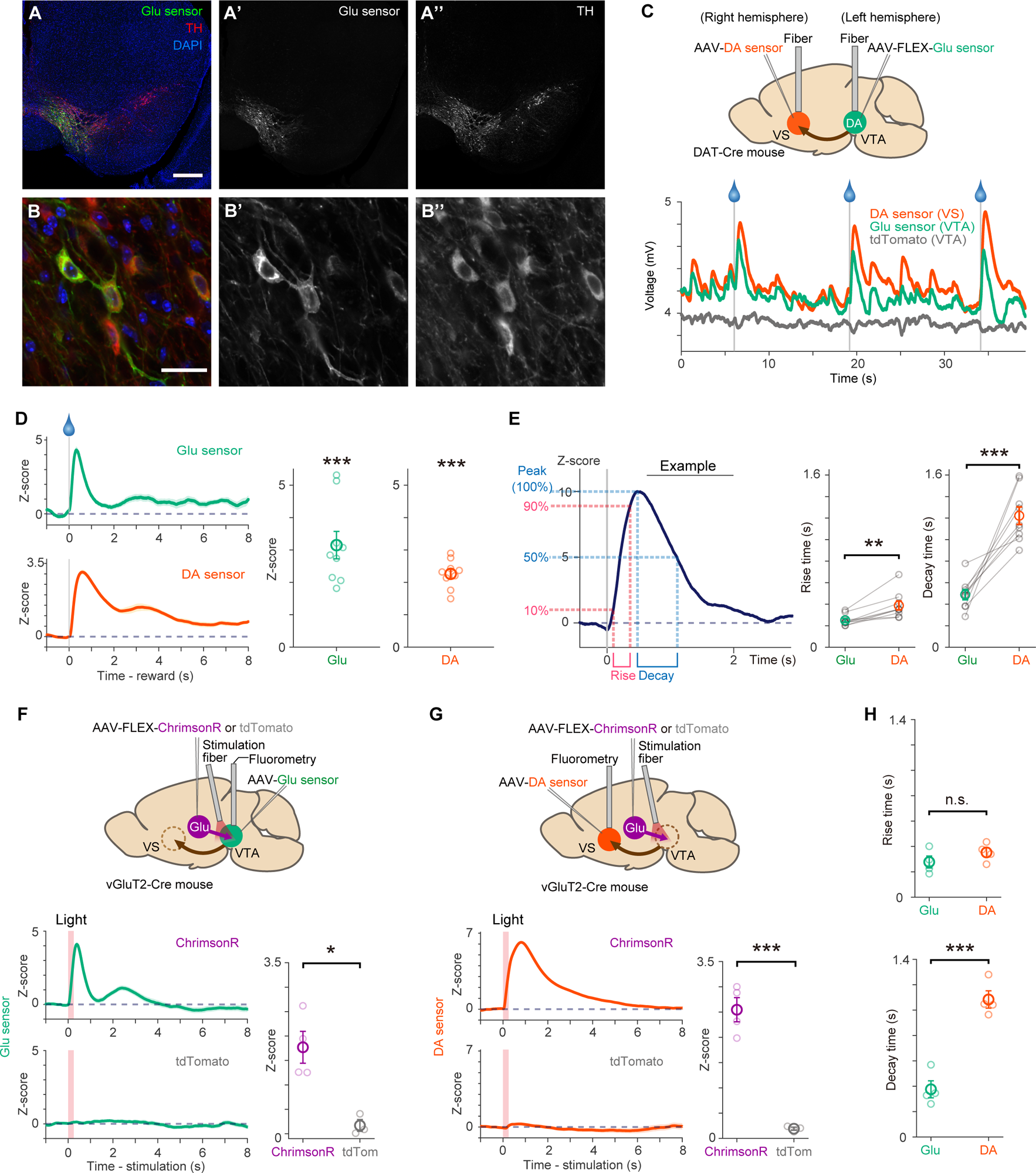
Characterization of glutamate sensor signals in the VTA. (A and B) Expression of glutamate sensor (SF-iGluSnFR) in the VTA (A) and its close-up view (B). Green, glutamate sensor; red, immunohistochemistry against tyrosine-hydroxylase (TH); blue, counter staining with DAPI. Scale bars, 500 μm (A) and 25 μm (B). (C) A schematic of the experimental design, and example raw signals for simultaneous recording of glutamate sensor signals in the VTA, control tdTomato in the VTA, and dopamine sensor signals in the VS. (D) Signals of glutamate sensor (left upper and middle; *t* = 7.4, *p* = 0.72×10^-4^, two-sided *t*-test) and dopamine sensor (left bottom and right; *t* = 15, *p* = 0.31×10^-6^, two-sided *t*-test) in response to unexpected water from an example session (left, mean ± s.e.m.) and each animal (n = 9 animals, middle and right, 1-500 ms from reward onset, mean ± s.e.m.). (E) Rise and decay time for glutamate sensor and dopamine sensor signals evoked by water reward (n = 9 animals). An example of rise and decay detection (left). Comparison of rise (middle; *t* = -4.6, *p* = 0.15×10^-2^, two-sided paired *t*-test) and decay (right; *t* = -6.5, *p* = 0.18×10^-3^, two-sided paired *t*-test) between sensors. Mean ± s.e.m. (F and G) Glutamate sensor signals at the VTA (F; *t* = 4.0, *p* = 0.010, two-sided unpaired *t*-test; n = 4 animals for ChrimsonR, n = 3 animals for tdTomato) and dopamine sensor signals at the VS (G; *t* = 8.2, *p* = 0.41×10^-3^, two-sided unpaired *t*-test; n = 4 animals for ChrimsonR, n = 3 animals for tdTomato) at optogenetic activation of glutamate axons (VP, PPTg, and STh) in the VTA from an example session (left, mean ± s.e.m.) and for each animal (right, 1-500 ms from light onset, mean ± s.e.m.). (H) Comparison of rise (upper; *t* = -1.2, *p* = 0.25, two-sided unpaired *t*-test) and decay times (bottom; *t* = -7.4, *p* = 0.31×10^-3^, two-sided unpaired *t*-test) for glutamate and dopamine sensor signals evoked by the optogenetic glutamate axon stimulation. Mean ± s.e.m.

To test causality between glutamate inputs and dopamine release, we optogenetically activated glutamate inputs to VTA. To do this, ChrimsonR^42^, a light-gated cation channel, was expressed in glutamate neurons in multiple pre-synaptic areas to dopamine neurons (pedunculopontine tegmental nucleus (PPTg), subthalamic nucleus (STh), and ventral pallidum (VP))^17,35^ in vesicular glutamate transporter 2 (vGluT2)-Cre mice, and their axons were activated at the VTA (Figures 1F and 1G). We first examined whether glutamate sensor signals reliably capture activation of glutamate inputs. We observed a significant increase of glutamate sensor signals when glutamate axons in VTA were stimulated in ChrimsonR-expressing animals, more than control mice expressing tdTomato (Figure 1F). The same optogenetic stimulation also strongly evoked dopamine release in the VS in ChrimsonR-expressing animals, more than in control mice (Figure 1G). We observed that glutamate sensor signals showed similar or faster responses to optogenetic stimulation (rise: 277±47.1 ms, decay: 375±66.9 ms) compared to dopamine release (rise: 352±36.7 ms, decay: 1083±68.1 ms) (Figure 1H). These observations confirmed that glutamate sensor recording with fiber-fluorometry allows detection of glutamate release in the VTA that causally evokes dopamine release in VS.

### Glutamate input sends RPE to dopamine neurons during classical conditioning

Multiple studies have proposed potential neural mechanisms for the computation of TD error signals in dopamine neurons^4,12–16,27^. Most models hypothesized that inhibitory inputs play key roles in TD error computation by providing reward expectation or prediction error that is already computed. According to these models, inhibitory input should be inhibited at reward cue to generate excitation in dopamine neurons. However, a previous study recorded neural activity from monosynaptic inputs to dopamine neurons and found that the vast majority of input neurons were activated by a reward cue^22^. This suggests that excitatory inputs to dopamine neurons, likely conveyed via glutamate, play critical roles in driving dopamine responses. Previous studies also found that glutamate inputs to the VTA are distributed throughout the brain^17^ and that presynaptic neurons signaling reward value are also distributed^22^. To probe the role of glutamate inputs in TD error calculations, we recorded population activity of glutamate input while mice performed a classical conditioning paradigm (Figure 2; Figure S1). We first examined whether glutamate input responses to reward-predicting cues are modulated by associated reward value, which is one of the characteristics of TD error (Figures 2B and 2D). We found that in response to odor cues, glutamate signals were monotonically modulated by the associated probabilities of reward outcome. To analyze the relationship between cue responses and associated reward, we performed linear regression of cue responses with associated reward probability for each animal. We observed a positive linear relationship between glutamate signals and reward probabilities (Figure 2E). While the observed activity patterns in glutamate input are consistent with TD error, this observation does not distinguish TD error from reward value.

**Figure 2.**
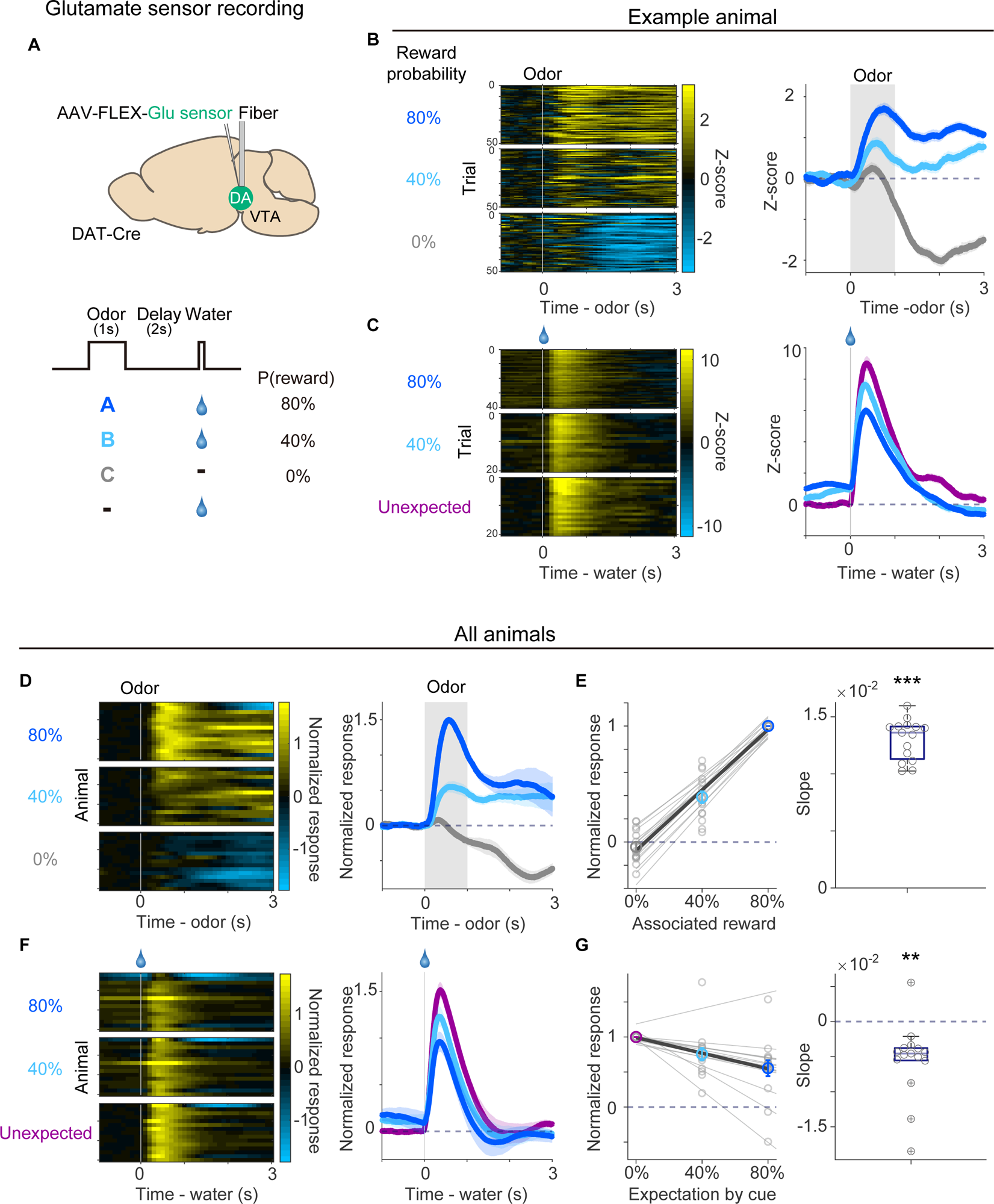
Recording of glutamate inputs to the VTA dopamine neurons during classical conditioning. (A) Schematics of recording of glutamate sensor signals from dopamine neurons and classical conditioning task. (B and C) Glutamate sensor responses to odor cues (B) and water reward (C) in each trial (left) and averaged activity of all trials (right) in a single session in an example animal. Mean ± s.e.m. (D) Average glutamate sensor responses to odor cues in each animal, normalized by 80% reward odor response (left) and averaged activity for all animals (right, n =15 animals). Mean ± s.e.m. (E) Linear regression of responses to odors (1-1000 ms from odor onset) with associated reward probability. Left, light gray, each animal; dark gray, average of all animals, correlation coefficient 0.94, *p* = 0.11×10^-21^, *F*-test. Right, regression coefficients for each animal (*t* = 28, *p* = 0.85×10^-13^, two-sided *t*-test, n =15 animals). (F) Average responses to water reward in each animal, normalized by unexpected reward response (left), and averaged activity for all animals (right). Mean ± s.e.m. (G) Linear regression of responses to water (201-1200 ms from reward onset) with expected reward. Left, light gray, each animal; dark gray, average of all mice, correlation coefficient - 0.71, *p* = 0.38×10^-7^, *F*-test. Right, regression coefficients for each animal (*t* = -3.9, *p* = 0.13×10^-2^, two-sided *t*-test). In box plots, grey lines are the median; edges are 25^th^ and 75^th^ percentiles; and whiskers are the most extreme data points not considered as outliers.

We next examined responses to water reward with different expectations (Figures 2C and 2F). The reward responses in glutamate inputs were negatively modulated by expectation; the higher the predicted reward probability was, the smaller the glutamate signal was (Figure 2G). The response patterns to water reward do not match with reward expectation or actual reward value. Rather, these results are more consistent with TD error coding where reward responses are suppressed when the reward is expected.

### Glutamate input activity follows TD error rules in sequential conditioning

A hallmark of TD error signal is that, in addition to rewarding outcomes, moment-by-moment changes in reward expectation (value) drives TD errors^11^. According to this model, a reduction of response occurs not only for a reward when that reward was predicted by a cue, but also for a reward-predicting cue when the cue was preceded by a different cue that itself predicts the upcoming reward (’sequential conditioning’, Figure 3A). We next sought to use this property to test for TD error coding in the glutamate input to dopamine neurons (Figure 3A right; Figure S1)^43,44^. Mice were first trained in simple classical conditioning to associate an odor (’proximal odor’) with either reward or no outcome (Figure 3A Step1). After learning the association, new odors (’distal odors’) were presented at an earlier time point (Figure 3A Step2). One distal odor was associated with reward-predicting cue at 100%, one with reward-predicting cue and no outcome cue at 50%, and one with no outcome cue at 100%. After completing the training, we simultaneously monitored glutamate input to dopamine neurons and dopamine release in the VS. As expected, dopamine release at the distal odor cue monotonically increased with the associated reward cue probability (Figures 3B and 3C). Critically, the magnitude of dopamine responses to the proximal reward-predicting cue (Odor A) was negatively scaled by the probability that this cue was predicted by a distal cue; strongest when there was no predicting cue, intermediate when this cue was predicted with 50% probability, and the smallest when the cue was predicted with 100% (Figures 3D and 3E). The dopamine activity pattern in this paradigm was consistent with TD errors in the TD learning model^11^.

**Figure 3.**
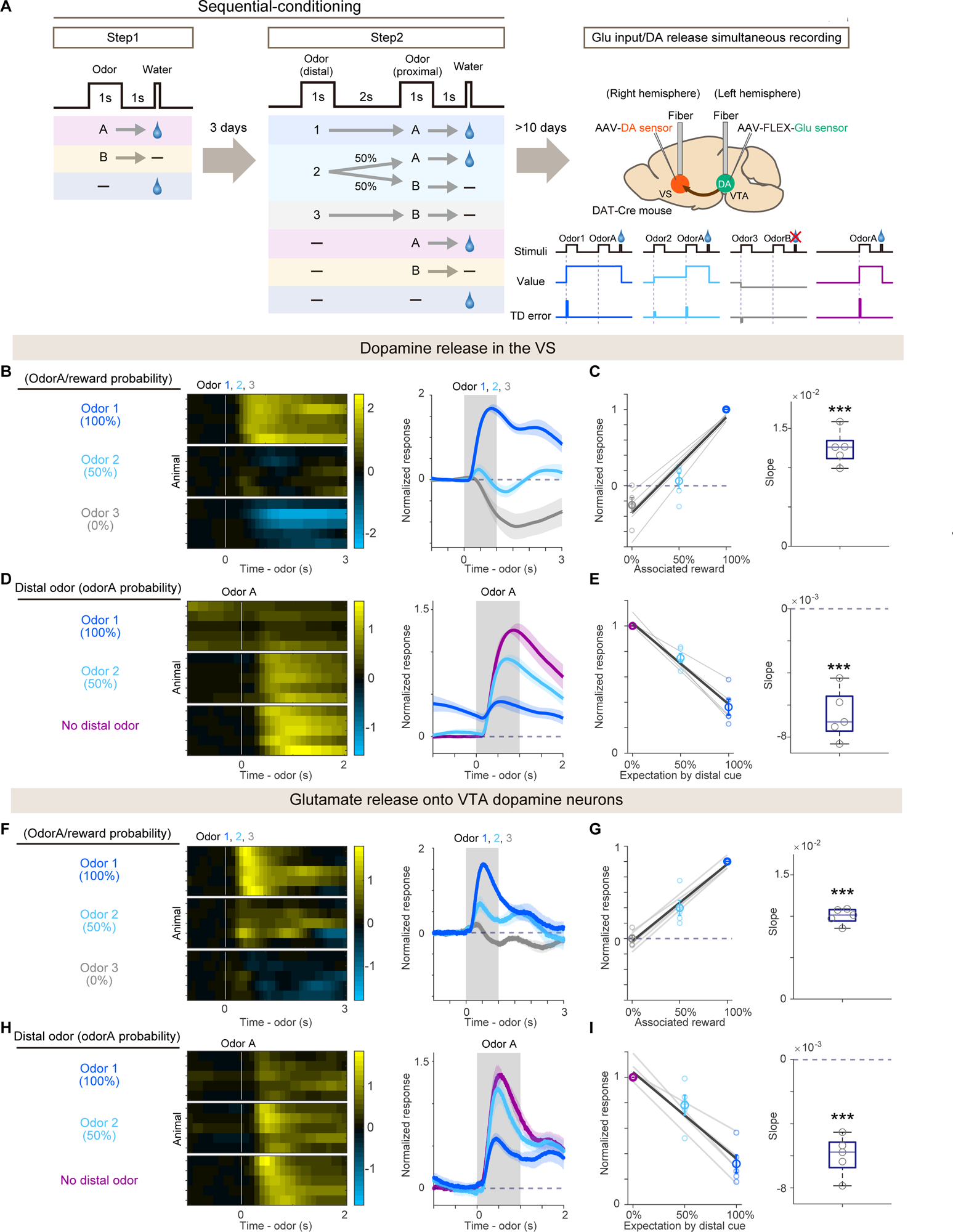
Simultaneously recorded glutamate sensor and dopamine sensor signals during sequential conditioning. (A) Schematics of sequential conditioning. The mice were trained to associate proximal cues and outcomes (Step1) and then distal odors were introduced to associate with proximal cues and outcomes (Step2). After completing sequential conditioning, glutamate sensor signals from the VTA dopamine neurons and dopamine sensor signal in the VS were recorded simultaneously (right top). Bottom right, expected value and TD error signals based on a TD learning model. (B) Dopamine responses to distal odor cues in each animal, normalized by odor 1 response (left), and averaged activity for all animals (right, n = 5 animals). Mean ± s.e.m. (C) Linear regression of responses to distal odors (1-1000 ms from distal odor onset) with associated reward probability. Left, light color, each animal; dark color, average of all mice, correlation coefficient 0.92, *p* = 0.11×10^-5^, *F*-test; mean ± s.e.m. Right, regression coefficients in each animal (*t* = 12, *p* = 0.20×10^-3^, two-sided *t*-test, n = 5 animals). (D) Dopamine responses to the reward-predicted proximal odor (odor A) in each animal, normalized with responses to unexpected odor A (left), and averaged activity for all animals (right, n = 5 animals). Mean ± s.e.m. (E) Linear regression of responses to odor A (201-1200 ms from odor A onset) with expectation of odor A. Left, light color, each animal; dark color, average of all mice, correlation coefficient - 0.94, *p* = 0.93×10^-7^, *F*-test; mean ± s.e.m.. Right, regression coefficients in each animal (right; *t* = -9.3, *p* = 0.72×10^-3^, two-sided *t*-test, n = 5 animals). (F) Responses of glutamate inputs to dopamine neurons to distal odor cue (n = 5 animals). Mean ± s.e.m. (G) Linear regression of responses at distal odors (1-1000 ms from distal odor onset) with associated reward probability. Left, light color, each animal; dark color, average of all mice, correlation coefficient 0.95, *p* = 0.44×10^-7^, *F*-test; mean ± s.e.m.. Right, regression coefficients in each animal (*t* = 23, *p* = 0.18×10^-4^, two-sided *t*-test). (H) Responses of glutamate inputs to dopamine neurons to odor A (n = 5 animals). Mean ± s.e.m. (I) Linear regression of responses to odor A (201-1200 ms from odor A onset) with expectation of odor A. Left, light color, each animal; dark color, average of all mice, correlation coefficient - 0.88, *p* = 0.12×10^-4^, *F*-test; mean ± s.e.m.. Right, regression coefficients in each animal (*t* = -10, *p* = 0.43×10^-3^, two-sided *t*-test, n = 5 animals). In box plots, grey lines are the median; edges are 25^th^ and 75^th^ percentiles; and whiskers are the most extreme data points not considered as outliers.

Like dopamine activity, glutamate release at the distal odors showed a monotonic increase with the probability of the reward cue (Figures 3F and 3G). Importantly, similar to dopamine release, the magnitude of glutamate signals at the proximal reward-predicting cue (Odor A) was negatively scaled by the probability that this cue was predicted by a distal cue (Figures 3H and 3I). Thus, glutamate input to dopamine neurons shows characteristics of TD error, resembling the observed patterns in dopamine release.

Notably, results from the sequential conditioning strengthen our observation in the simple classical conditioning in two ways. First, because cue events are temporally separated from actual reward acquisition, potential movement-related neural activity or recording noise caused by anticipatory licking were greatly reduced (Figure S2). We observed similar activity patterns both in unprocessed glutamate sensor signals and in normalized signals with control fluorescent signals (Figure S2; see Methods). Second, expectation-dependent reduction of cue responses in addition to water responses generalizes the idea of RPE to TD error, in which the value of the proximal cue takes the place of reward when computing the difference in value expectations across time. Our observations thus indicate that both dopamine release and glutamate inputs comply with TD error rules (i.e. computing changes in values) in sequential conditioning tasks.

### Glutamate input shows inhibition at omission of expected outcome

Omission of expected reward produces dopamine inhibition below baseline^4,31^. The inhibitory “dip” at reward omission is another characteristic of TD error, because it can be explained by a prediction of reward that fails to materialize. It is commonly assumed that the inhibitory responses to reward omission are driven solely by GABA inputs such as GABA neurons in the RMTg^26^. However, we found that glutamate input as well as dopamine activity and release showed significant inhibition when an expected reward was omitted during classical conditioning (Figures 4A and 4B; Figure S1). Similarly, TD learning predicts a negative prediction error at the reward omission cue (a cue signaling no outcome when reward cue has been expected) in sequential conditioning. Consistently, dopamine activity dipped below baseline when the reward omission cue appeared in sequential conditioning (Figures 4C-4E). Notably, glutamate input activity also showed an inhibitory dip at the reward omission cue (Figures 4D and 4E).

**Figure 4.**
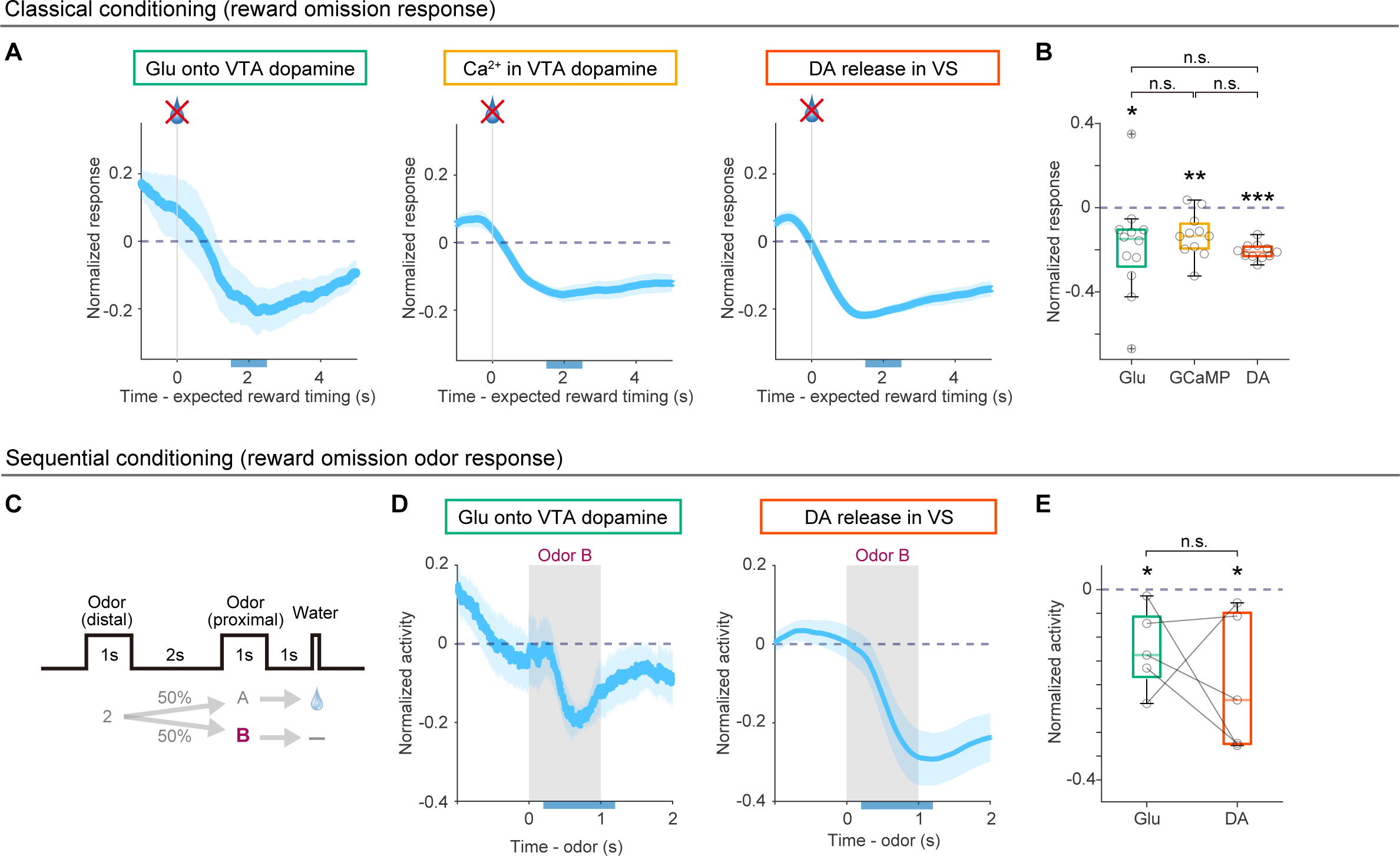
Responses to reward omission in glutamate inputs to dopamine neurons, dopamine cell body activity and dopamine release. (A) Responses to omission of 40% expected reward in glutamate sensor in dopamine neurons in the VTA (left, n = 12 animals), GCaMP signal at dopamine cell bodies in the VTA (middle, n = 11 animals), and dopamine sensor signal in the VS (right, n = 11 animals) normalized with responses to unexpected water reward. Mean ± s.e.m. (B) Quantification of omission responses (1501-2500 ms from expected reward timing, Glu, *t* = -2.6, *p* = 0.022; GCaMP, *t* = -4.1, *p* = 0.18×10^-2^; DA, *t* = -18, *p* = 0.53×10^-8^; two-sided *t*-test) and comparison of different sensor signals (*F* = 0.70, *p* = 0.50, one-way ANOVA; Glu vs GCaMP, *p* = 0.69, Glu vs DA, *p* = 0.92, GCaMP vs DA, *p* = 0.48, Tukey’s test). (C) Schematics of a proximal odor (odor B) that signals reward omission after a distal odor (odor 2) that signals 50% reward in sequential conditioning. (D) Responses to a reward omission odor in simultaneously recorded glutamate sensor signals (left) and dopamine sensor signals (right). n = 5 animals. Mean ± s.e.m. (E) Quantification of responses to a reward omission odor in glutamate sensor (201-1200 ms from odor B onset, *t* = -3.2, *p* = 0.032, two-sided *t*-test) and dopamine sensor (*t* = -3.0, *p* = 0.039, two-sided *t*-test), and comparison of these signal (*t* = -0.77, *p* = 0.48, two-sided paired *t*-test). In box plots, lighter color lines are the median; edges are 25^th^ and 75^th^ percentiles; and whiskers are the most extreme data points not considered as outliers.

We observed that glutamate input activity shows multiple characteristics of TD error in classical conditioning; cue responses were increased by associated reward value, reward responses were decreased by reward expectation, and reward omission induced inhibitory responses. As TD learning predicts, expectation-dependent suppression of responses was not restricted to reward responses. In sequential conditioning, we observed that reward cue responses were suppressed by expectation, and that the reward omission cue induced inhibitory responses. Together, these observations indicate that glutamate inputs to dopamine neurons convey significantly more complete TD error than popular theories predict^12–14,25,45^.

### TD errors in glutamate inputs are positively biased compared to dopamine cell body activity and dopamine release

Because we observed striking similarities between glutamate inputs to dopamine neurons in the VTA and dopamine release in VS, we directly compared their activity patterns. We first focused on signals in sequential conditioning recorded simultaneously. While both glutamate inputs and dopamine release showed characteristics of TD error, the temporal patterns were slightly different. Because glutamate sensor signals in the VTA and dopamine sensor signals in the VS show different temporal patterns of responses to the same optogenetic stimulation (Figures 1F-1H), we first estimated dopamine sensor signals in sequential conditioning solely by glutamate sensor signals. To do that we deconvolved glutamate signals using “glutamate kernels” estimated from the optogenetic responses in glutamate sensor signals, and then convolved the resulting trace with “dopamine kernels” estimated from the dopamine sensor optogenetic responses (Figures 5A-5C; see Methods^46^). With this method, glutamate input signals explained 54±19 % of variance in dopamine signals in sequential conditioning (Figures 5A-5E). While the estimated glutamate input contribution in dopamine distal odor responses were monotonically increased by associated reward odor probability (glutamate: *t* = 3.0, *p* = 0.038, *t*-test; n = 5 animals; Figure 5F left and right), the residual signal did not show significant modulation (residual: *t* = 0.47, *p* = 0.66, *t*-test; n = 5 animals, glutamate vs residual: *t* = 3.6, *p* = 0.022, paired *t*-test; n = 5 animals; Figure 5F middle and right). Similarly, the glutamate contribution, but not residual, at the proximal reward-predicting cue (Odor A) was negatively scaled by the probability that this cue was predicted by a distal cue (glutamate: *t* = -5.59, *p* = 0.0050, *t*-test; n = 5 animals; Figure 5G left and right; residual: *t* = -1.3, *p* = 0.25, *t*-test; n = 5 animals, glutamate vs residual: *t* = -4.3, *p* = 0.012, paired *t*-test; n = 5 animals; Figure 5G middle and right). The lack of TD error characteristics in residual signals suggests that glutamate input explains a significant portion of TD error coding in dopamine activity. While we observed striking similarities in estimated and actual dopamine signals, we also noticed differences. The cue responses predicted from the observed glutamate signals tended to be more positive, while actual dopamine cue responses were more negative, with the no-outcome cue generating inhibitory dopamine responses (Figures 5B, 5D, and 5E).

**Figure 5.**
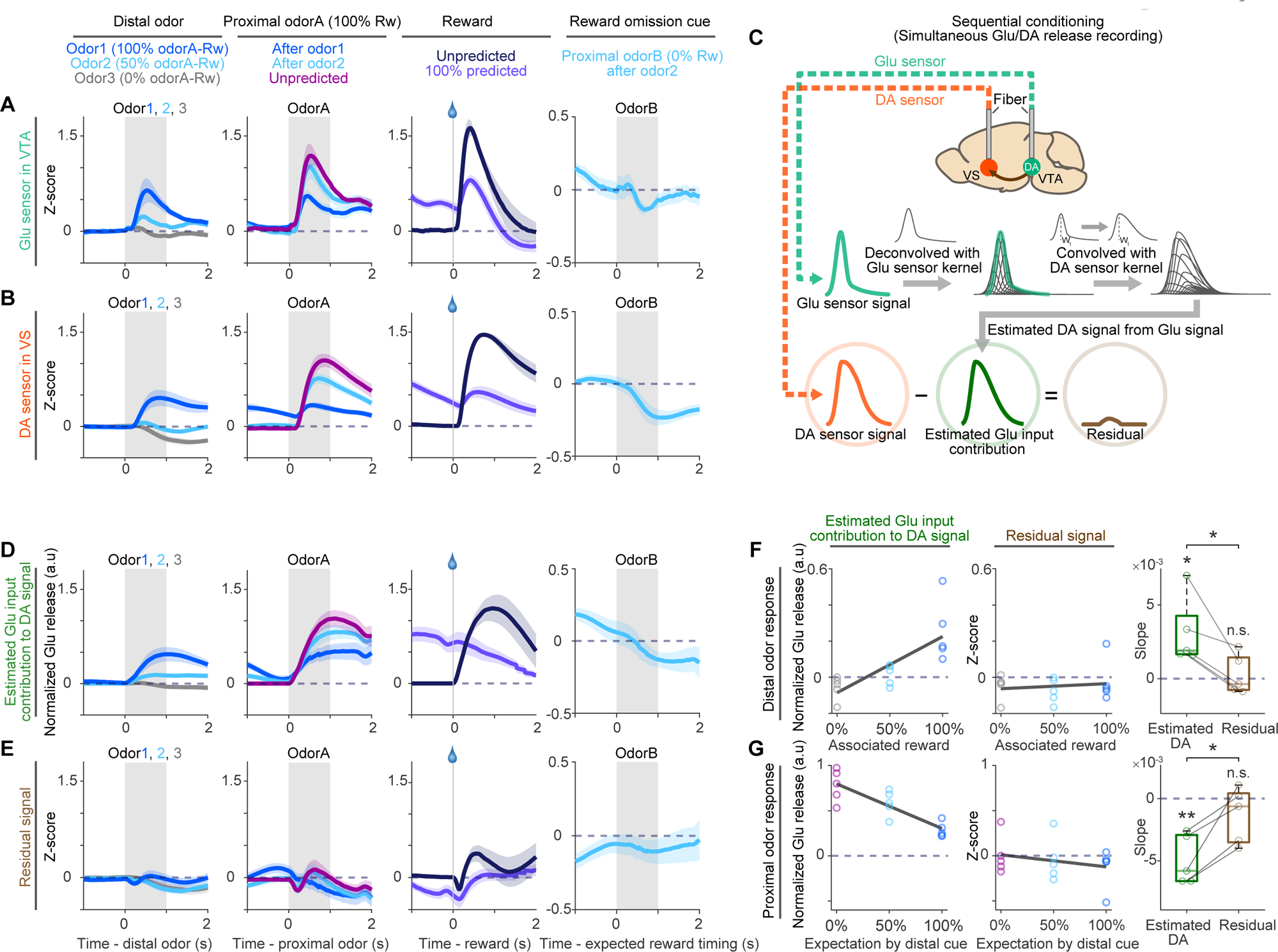
Estimation of dopamine signals from glutamate inputs in sequential conditioning. (A) Glutamate sensor signals at distal odors associated with different reward probability, a reward-predicted proximal odor with different expectation of the odor, water reward with different expectation, and a proximal reward omission odor (ordered left to right). Mean ± s.e.m. (B) Simultaneously recorded dopamine sensor signals. Mean ± s.e.m. (C) Dopamine signals were estimated by transforming glutamate sensor signals using differences in response patterns of glutamate sensor signals and dopamine sensor signals by optogenetic activation of glutamate inputs. (D) Estimated glutamate input contribution to dopamine sensor signals. Mean ± s.e.m. (E) Residual dopamine sensor signals that are not explained by glutamate sensor signals. Mean ± s.e.m. (F and G) Linear regression of responses to distal odors (F, 1-1000 ms from distal odor onset) or a proximal “odor A” (G, 201-1200 ms from odor A onset) with associated reward probability. Estimated glutamate contribution (distal odors: correlation coefficient 0.76, *p* = 0.89×10^-3^, *F*-test; proximal odors: correlation coefficient -0.84, *p* = 0.69×10^-4^, *F*-test; left), residual signal (distal odors: correlation coefficient 0.14, *p* = 0.61, *F*-test; proximal odors: correlation coefficient -0.25, *p* = 0.35, *F*-test; middle), and comparison of coefficient beta (right). n = 5 animals. In boxplots, lighter color lines are the median; edges are 25^th^ and 75^th^ percentiles; and whiskers are the most extreme data points not considered as outliers.

We next compared the activity pattern at each step of neural transmission: glutamate input to dopamine neurons in VTA (glutamate sensor in dopamine neurons), somatic calcium signals in VTA dopamine neurons (GCaMP in dopamine neurons), and dopamine release in the VS (dopamine sensor in striatal neurons) during classical conditioning (Figures 6A and 6B; Figure S1). While cue responses were monotonically modulated by associated reward value in all steps, we noticed that glutamate input responses were biased toward excitation. Glutamate input responses to 40% reward odor and 0% reward odor were significantly higher compared to dopamine somatic calcium signals and dopamine release (Figure 6C). We next estimated the minimum associated reward probability for the cue to produce positive responses (“zero-crossing point”; Figure 6D left) by linearly fitting neuronal responses with reward probability. The zero-crossing points were significantly lower in glutamate input activity (4.3±3.5 %) compared to dopamine cell body activity (32±3.0 %) and dopamine release (28±2.2 %) (Figure 6D right). This indicates that glutamate input responses to cues are positively biased compared to dopamine neuron activity. To test for a possible contribution of recording sites to the observed difference, we recorded glutamate and calcium sensor signals from varied locations along the mediolateral axis of the VTA (0.325-0.75 mm from the midline). We found that the difference between glutamate input activity and dopamine neuron activity in 40% reward odor response, 0% reward odor response, or zero-crossing point was not explained by the recording location (Figure 6E).

**Figure 6.**
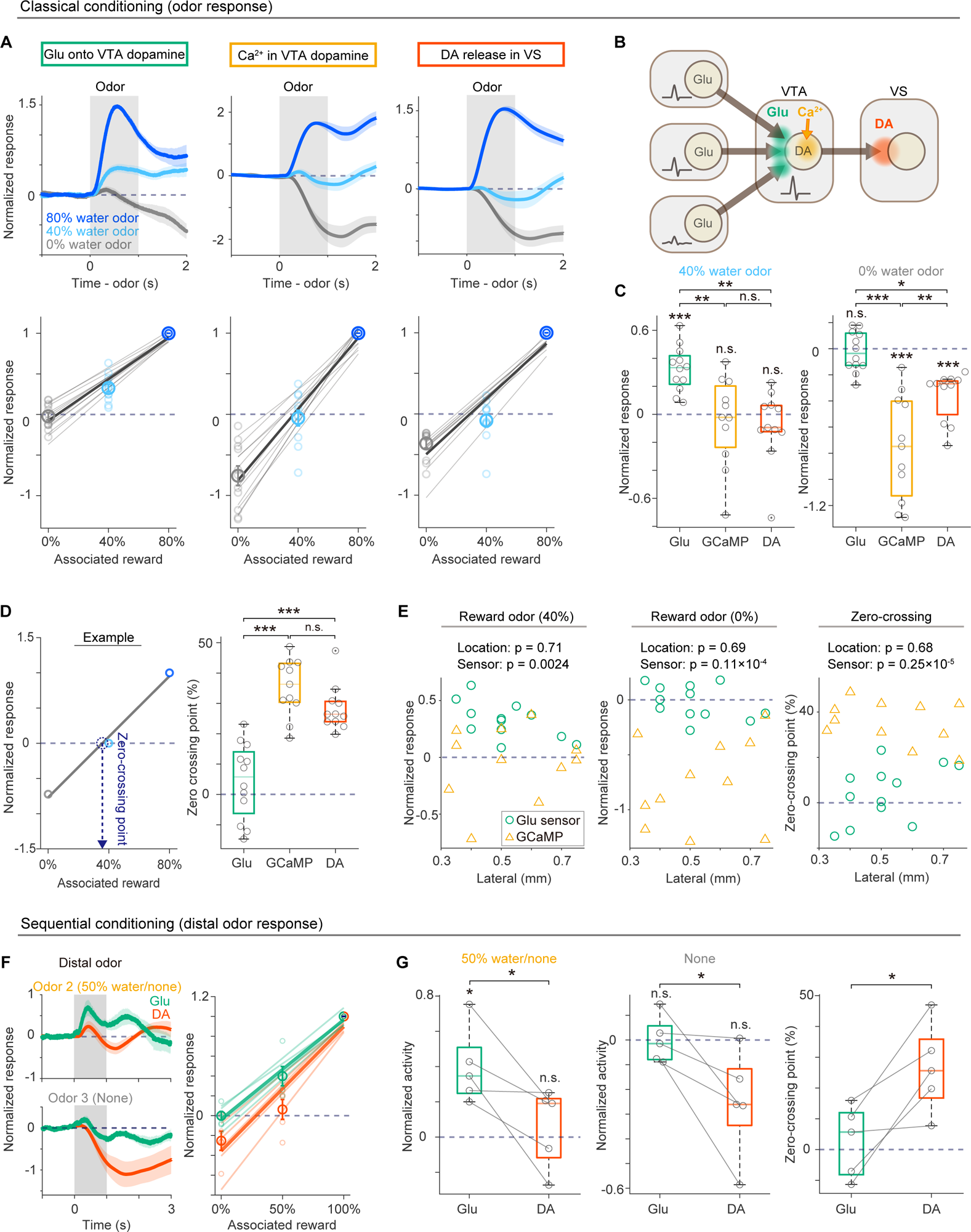
Comparison of glutamate input activity to dopamine neurons with dopamine cell body activity and dopamine release. (A) Responses to odors associated with different reward probability in classical conditioning. Left, glutamate sensor signal from the VTA dopamine neurons (n = 12 animals). Middle, GCaMP signals from dopamine cell bodies in VTA (n = 11 animals). Right, dopamine release in the VS (n = 11 animals). Mean ± s.e.m. Bottom, 1-1000 ms from odor onset. (B) Schematics of three different events during dopamine neural transmission. (C) Comparison of responses to 40% reward-predicted odor (1-1000 ms from odor onset, left; *F* = 9.2, *p* = 0.69×10^-3^, one-way ANOVA; Glu vs GCaMP, *p* = 0.37×10^-2^, Glu vs DA, *p* = 0.15×10^-2^, GCaMP vs DA, *p* = 0.94, Tukey’s test; Glu, *t* = 6.9, *p* = 0.23×10^-4^, GCaMP, *t* = -0.48, *p* = 0.63, DA, *t* = 1.0, *p* = 0.32, two-sided *t*-test), and an odor that is associated with no outcome (right; *F* = 21, *p* = 0.12×10^-5^, one-way ANOVA; Glu vs GCaMP, *p* = 0.68×10^-6^, Glu vs DA, *p* = 0.012, GCaMP vs DA, *p* = 0.45×10^-2^, Tukey’s test; Glu, *t* = -0.5, *p* = 0.62, GCaMP, *t* = -6.2, *p* = 0.96×10^-4^, DA, *t* = -6.5, *p* = 0.62×10^-4^, two-sided *t*-test). (D) Comparison of zero-crossing point (i.e. estimated reward probability when responses to a cue become zero; *F* = 29, *p* = 0.58×10^-7^, one-way ANOVA; Glu vs GCaMP, *p* = 0.89×10^-7^, Glu vs DA, *p* = 0.92×10^-5^, GCaMP vs DA, *p* = 0.25, Tukey’s test). (E) Effect of recording sites and sensors in responses to 40% reward-predicted odor (left; location, *F* = 0.14, *p* = 0.71; sensor, *F* = 12, *p* = 0.24×10^-2^, two-way ANOVA), responses to an odor predicting no outcome (middle; location, *F* = 0.15, *p* = 0.69; sensor, *F* = 33, *p* = 0.11×10^-4^, two-way ANOVA), and zero-crossing points (right; location, *F* = 0.17, *p* = 0.68; sensor, *F* = 42, *p* = 0.25×10^-5^, two-way ANOVA). (F) Comparison of responses to distal odors associated with different reward probability (1-1000 ms from distal odor onset). Mean ± s.e.m. n = 5 animals. (G) Comparison between glutamate and dopamine sensor signals for responses to a distal odor that predicts 50% reward-predicted proximal odor (left; 1-1000 ms from distal odor onset *, t* = 3.0, *p* = 0.038, two-sided paired *t*-test), responses to a distal odor that predicts an odor with no outcome (middle; 1-1000 ms from distal odor onset*, t* = 2.7, *p* = 0.049, two-sided paired *t*-test), and zero-crossing points of distal odor response (right; *t* = -3.5, *p* = 0.038, two-sided paired *t*-test). In boxplots, lighter color lines are the median; edges are 25^th^ and 75^th^ percentiles; and whiskers are the most extreme data points not considered as outliers.

We next confirmed that the observed difference was not due to variability across animals. We compared simultaneously recorded glutamate input activity and dopamine release during sequential conditioning. As seen during the odor response in classical conditioning, responses to distal odors were positively biased in glutamate inputs compared to dopamine release in single animals (50% reward distal odor: *t* = -3.0, *p* = 0.038, paired *t*-test; 0% reward distal odor: *t* = - 2.7, *p* = 0.049, paired *t*-test; zero-crossing point: Glu, 2.7±5.1 %, DA, 26±6.5 %, *t* = 3.5, *p* = 0.023, paired *t*-test, Figures 6F and 6G). Thus, the population activity of glutamate inputs does not fully explain dopamine activity, suggesting that dopamine neurons do not purely relay information from glutamate input population but require specific inputs, probably additional inputs, to produce more negative responses.

### Lack of inhibitory responses to aversive stimuli in glutamate inputs to dopamine neurons

Many dopamine neurons are inhibited by aversive stimuli such as air puff to the eye, consistent with the negative value of such stimuli^7,24,31^. We confirmed that both dopamine neuron activity in the VTA and dopamine release at the VS showed activation to water reward and inhibition to air puff (Figures 7A-7C; Figure S3). Surprisingly, however, while glutamate input showed activation to water reward, it was also phasically activated by the air puff (Figures 7A-7C; Figure S3A). Thus, the glutamate input response to the air puff is clearly different from dopamine activity or release in this task condition (Figure 7C). The difference was not explained by recording location (Figure 7D). The opposite directions in air puff response between dopamine activity and glutamate inputs strongly suggest requirement of an additional input, such as RMTg GABA neurons^25,27^, to cancel out excitation and produce inhibitory responses to aversive air puffs in dopamine neurons. In contrast to common models in which glutamate and GABA inputs provide components of TD errors (Figure 7E) or dopamine neurons inherit already calculated RPE from GABA neurons (Figure 7F), our results indicate that glutamate inputs send TD errors, including inhibitory responses to reward omission, but diverge from dopamine neurons in their response to aversive stimuli, showing excitation rather than inhibition. This excitation may compete with GABA inputs, which are also excited by aversive stimuli (Figure 7G).

**Figure 7.**
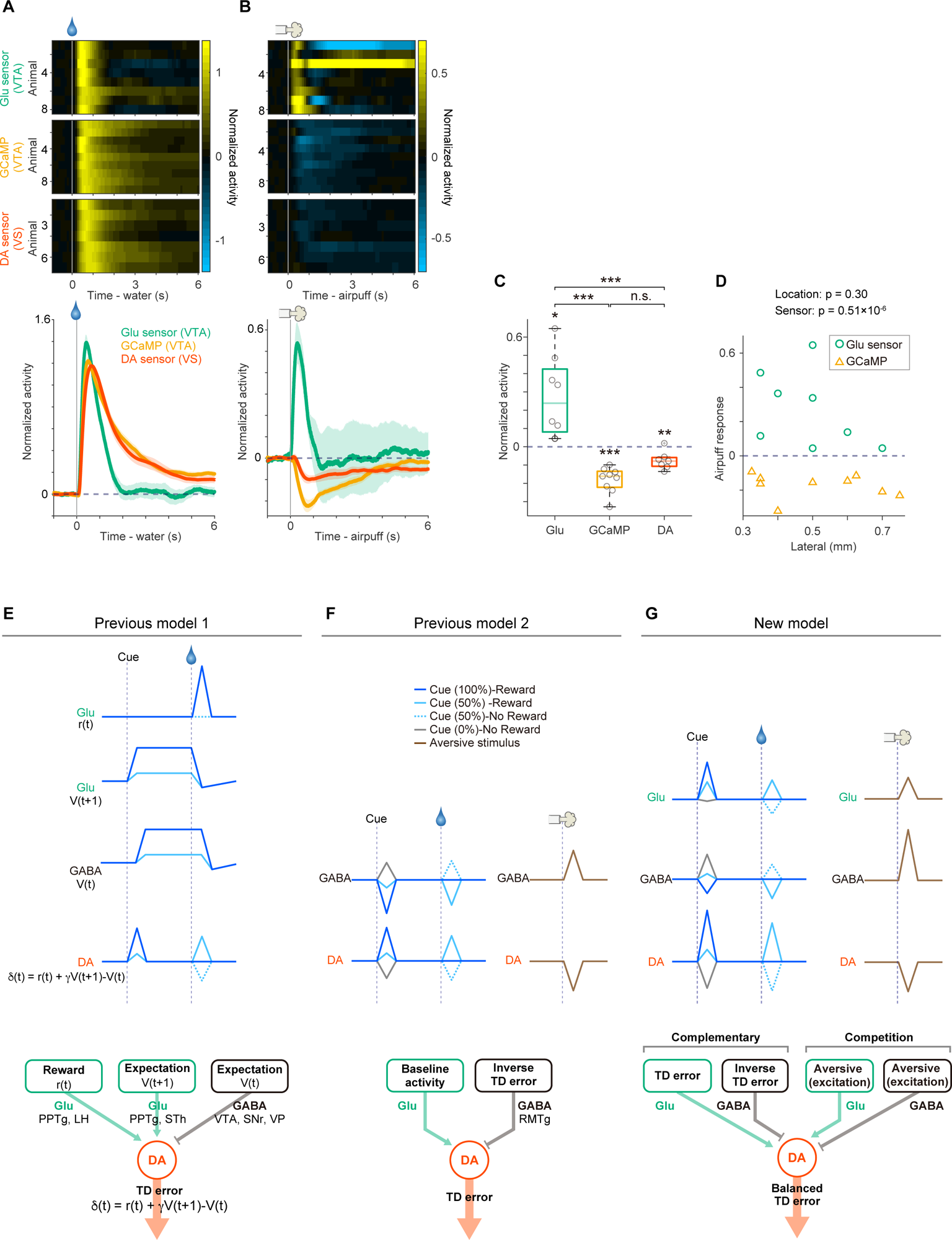
Responses to air puff in glutamate input activity to dopamine neurons, dopamine cell body activity and dopamine release. (A) Responses to unexpected water reward. (B) Responses to unexpected air puff. (C) Average responses to air puff (201-1200 ms from air puff onset, Glu; *t* = 3.4, *p* = 0.010, n = 8 animals, GCaMP; *t* = -7.6, *p* = 0.58×10^-4^, n = 9 animals, DA; *t* = -4.0, *p* = 0.69×10^-2^, n = 7 animals, two-sided *t*-test) and comparison of these responses (*F* = 24, *p* = 0.33×10^-5^, one-way ANOVA; Glu vs GCaMP, *p* = 0.22×10^-3^, Glu vs DA, *p* = 0.31×10^-5^, GCaMP vs DA, *p* = 0.30, Tukey’s test). Mean ± s.e.m. In boxplots, lighter color lines are the median; edges are 25^th^ and 75^th^ percentiles; and whiskers are the most extreme data points not considered as outliers. (D) Effects of recording sites and sensors in air puff responses (location, *F* = 1.1, *p* = 0.30; sensor, *F* = 32, *p* = 0.51×10^-6^, two-way ANOVA). (E) Classical models for TD error computation in dopamine neurons. Different components for TD error computation are conveyed by input to dopamine neuron; GABA input conveys expectation (V(t)), and glutamate input conveys expectation (V(t+1)) and reward (r(t))^12,14^ (alternatively expectation (V(t+1)) is provided by GABA input as inverse form^12,15^). γ; temporal discounting factor (0 ≤ γ ≤1). (F) Disinhibition model. RMTg GABA inputs provide TD error through disinhibition of dopamine neuron^25,27^. Glutamate input simply provides baseline activity^61^. (G) Schematics of dopamine signal computation. Glutamate input from various areas conveys positively biased TD error as a population. GABA input from RMTg (and potentially other areas) also sends TD error. Combination of these redundant inputs forms balanced TD error in dopamine neurons. Negative value of aversive events is conveyed by GABA input, whereas glutamate input is excited by aversive events, opposing to GABA inputs.

### Opioids may flip dopamine responses to aversive stimuli from inhibition to excitation

If both glutamate inputs and GABA inputs are excited by aversive stimuli, and thus compete each other to shape dopamine responses, the dopamine responses to aversive stimuli may be flexibly altered by slight modulations in the relative strength of excitatory and inhibitory inputs, depending on an animal’s state or context (Figure 7G). We focus here on one state alteration: the exogenous administration of opioids, common and effective analgesics which lead to an increase in dopamine excitability and inhibit some direct and indirect inputs to dopamine neurons^47–54^. We tested the effect of systemic administration of buprenorphine, an opioid commonly used in medical practice, on dopamine release and glutamate inputs with simultaneous recording (Figure 8; Figure S3). While dopamine release in VS was inhibited at an aversive air puff in the control sessions, buprenorphine treatment drastically diminished inhibition of the dopamine release and even flipped the response to activation in some animals (Figures 8A-8B). The reduction of dopamine inhibition was specifically observed in buprenorphine treatment while control saline injection did not alter dopamine response to air puff (Figure 8B right). In contrast, glutamate input responses were not clearly changed by the buprenorphine treatment (Figures 8C-8D). Neither buprenorphine nor saline administration altered excitatory responses in glutamate input (Figure 8D right). Overall, buprenorphine administration consistently decreased the magnitude of dopamine inhibitory response to an aversive stimulus, a change which was not seen in glutamate inputs (Figure 8E), suggesting competition between glutamate inputs and other inputs. In this way, excitation by aversive stimuli in glutamate and other inputs appears to be differentially modulated to flexibly shape dopamine responses to aversive stimuli.

**Figure 8.**
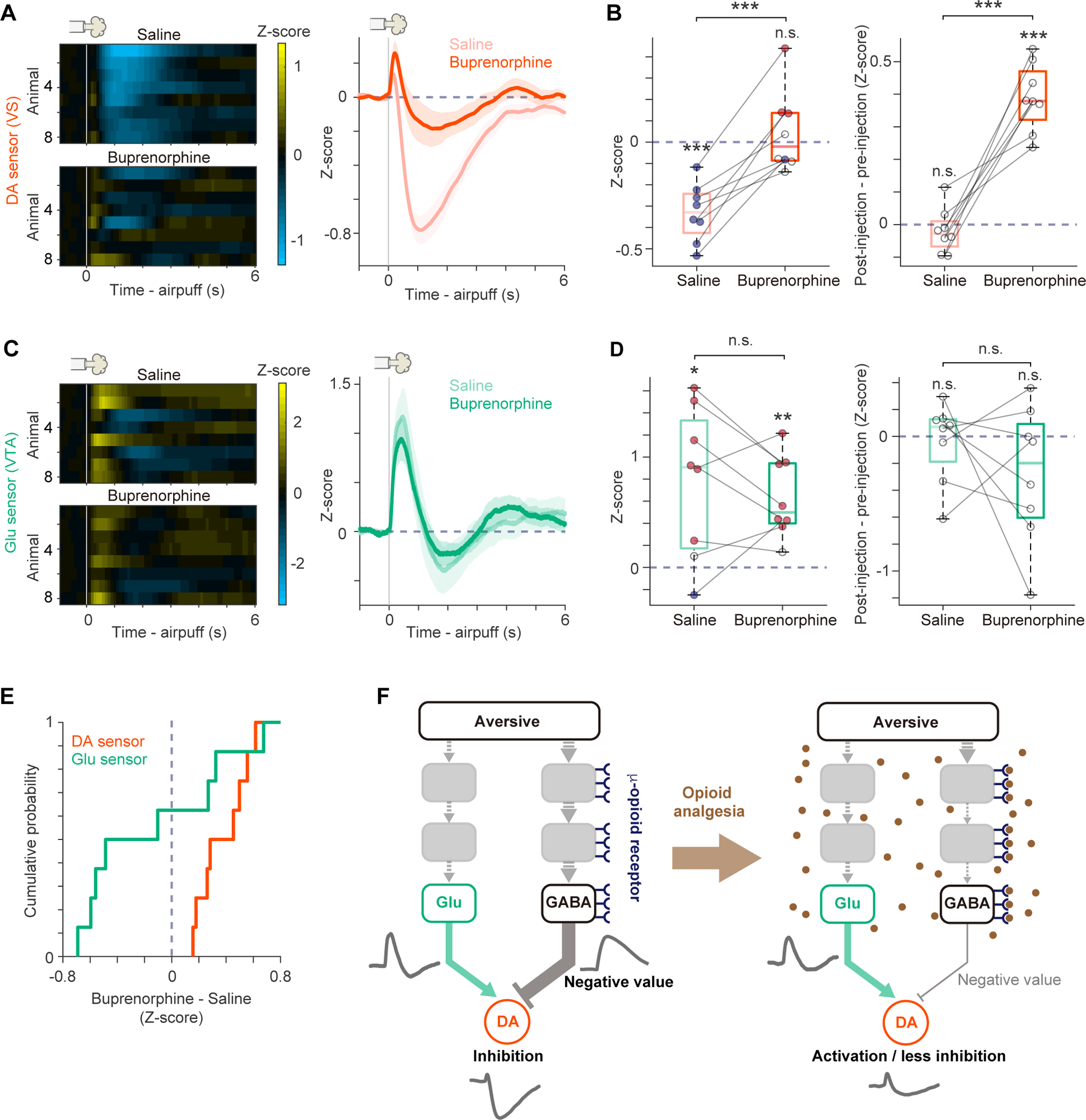
Effect of buprenorphine on dopamine activity and glutamate input in response to aversive air puff. (A) Dopamine release in the VS at unexpected air puff after saline or buprenorphine injection. Mean ± s.e.m. (B) Left, comparison of dopamine release following air puff after saline and buprenorphine injection (1-1000 ms from air puff onset, saline vs buprenorphine: *t* = 5.9, *p* = 0.55×10^-3^, paired t-test, saline: *t* = -6.9, *p* = 0.23×10^-3^, *t*-test, buprenorphine: *t* = 0.66, *p* = 0.52, *t*-test, n = 8 animals). Right, difference in response to air puff between pre-injection and post-injection (1-1000 ms from air puff onset, saline vs buprenorphine: *t* = 7.2, *p* = 0.16×10^-3^, paired *t*-test, saline: *t* = -0.78, *p* = 0.46, *t*-test, buprenorphine: *t* = 10, *p* = 0.14×10^-4^, *t*-test, n = 8 animals). A circle indicates each animal. Blue circle: significantly below 0 (0.05 < *p*, 0 < *t*, *t*-test). Red circle: significantly above 0 (0.05 < *p*, 0 > *t*, *t*-test). (C) Glutamate input to dopamine at air puff after saline or buprenorphine injection. Mean ± s.e.m. (D) Left, comparison of glutamate input response to air puff after saline and buprenorphine injection (1-1000 ms from air puff onset, saline vs buprenorphine: *t* = -0.79, *p* = 0.45, paired *t*-test, saline: *t* = 3.2, *p* = 0.014, *t*-test, buprenorphine: *t* = 4.8, *p* = 0.17×10^-2^, *t*-test, n = 8 animals). Right, difference in response to air puff between pre-injection and post-injection (1-1000 ms from air puff onset, saline vs buprenorphine: *t* = -1.0, *p* = 0.33, paired *t*-test, saline: *t* = -0.35, *p* = 0.73, *t*-test, buprenorphine: *t* = -1.5, *p* = 0.16, *t*-test, n = 8 animals). Circles indicate individual animals. Blue circle: significantly below 0 (0.05 < *p*, 0 < *t,* t-test). Red circle: significantly above 0 (0.05 < *p*, 0 > *t,* t-test). (E) Distribution of changes in dopamine and glutamate input response to air puff with buprenorphine and saline injection (*ks* = 0.62, *p* = 0.049, Kolmogorov-Smirnov test, n = 8 animals). (F) Potential competition between glutamate and GABA inputs to dopamine neurons. At an aversive event, a large increase of inhibitory inputs cancels out the increase of excitatory inputs and generates an inhibitory dip in dopamine activity. With opioid administration, dopamine shows more activation and less inhibition in response to aversive events, potentially because of preferential inhibition of GABA inputs by opioids. In boxplots, lighter color lines are the median; edges are 25^th^ and 75^th^ percentiles; and whiskers are the most extreme data points not considered as outliers.

## Discussion

Excitatory-Inhibitory balance has been intensively studied in the context of neural dynamics, sensory physiology and psychiatric conditions^55–59^, and it is a common idea that inhibitory tones should be suitably titrated for specific roles such as stabilizing feedback loops or sharpening the stimulus selectivity of sensory responses^60^. Our study points out a critical similarity and difference between dopamine activity and its excitatory inputs activity. Together with reported GABA input activity^25–27,31^, our finding suggests a different aspect of excitatory-inhibitory balance: both glutamate and GABA inputs provide overlapping TD error characteristics to dopamine neurons, while they compete with each other for control over dopamine responses to aversive stimuli. These two contrasting findings suggest that slight modifications of excitatory-inhibitory weights of inputs would dramatically flip responses to aversive stimuli but preserve core information of TD errors in dopamine neurons.

We examined information conveyed by glutamate inputs to dopamine neurons for RPE computation. The prevailing view of RPE computation in dopamine neurons is that inhibitory neurons are the major contributor of prediction error signals in dopamine neurons. For example, because of the heavy projection from inhibitory inputs to dopamine neurons (>70 % of all inputs)^20^, inhibitory neurons are often hypothesized to provide prediction error signals to dopamine neurons via disinhibition (Figure 7F)^12,13,15,16,25,27,61^. Recent findings showed that inhibitory projections from the lateral hypothalamus (LH), VS, and superior colliculus (SC) to VTA GABA neurons are capable of disinhibiting dopamine neuron^18,37,38^, although optogenetically identified GABA neurons in VTA do not show inhibitory responses to reward or reward-predicting cues^31^. In simple RPE models, it has been proposed that inhibitory inputs provide expectation and excitatory inputs provide actual reward information^12^. Further, in models of TD learning, excitatory and inhibitory inputs both provide reward expectation (or value) but at slightly shifted timing so that dopamine neurons perform subtraction (or differentiation) to compute TD errors, a specific form of RPE (Figure 7E)^12,14^. In stark contrast to these views, we observed striking similarity between glutamate input activity and TD error itself. Notably, glutamate signals are negatively modulated by prior expectation, opposite to the idea that they encode expectation signals (Figure 7G).

A previous study found that information for reward expectation (value) and reward is distributed and mixed in single presynaptic neurons to dopamine neurons across various brain areas^22^. Interestingly, they found that the simple summation of those input activities shows characteristics of RPE. These results indicated that inputs with mixed information are sufficient to produce RPE. However, because the previous study could not distinguish the cell types of recorded neurons (i.e. which neurotransmitters they release), the summation did not reflect actual circuit operations. Here, we extended this investigation to show that the population activity of glutamate inputs to dopamine neurons, which are likely distributed across brain areas^17^, encodes TD error.

Although the activity patterns of glutamate inputs strikingly resembled those of dopamine neurons, we also found differences between their activities. Overall, glutamate input responses were positively biased compared to dopamine responses. Another important difference is that, although glutamate input showed an inhibitory dip by reward omission, similar to dopamine, glutamate input was *activated* by aversive air puff, in contrast to dopamine, which shows inhibitory responses (Figure 7). These results suggest that an additional input is required to generate inhibitory responses to aversive air puffs in dopamine activity. Consistently, GABA inputs to dopamine neurons such as the lHb-RMTg pathway^21,25–27,62^ and GABA neurons in the VTA^31^ are activated by aversive stimuli, and thus may send negative value information to dopamine neurons.

In addition to providing information about aversive events, the lHb-RMTg pathway is thought to provide inhibitory responses at reward omission in dopamine neurons (Figure 7F). Ablation of either Hb or RMTg specifically impairs omission responses of dopamine neurons but largely preserves other responses such as reward cue responses^24,26^. Although these results indicate an important role of this pathway in reward omission responses in dopamine neurons, we found that glutamate input also responds to reward omission with an inhibitory dip. Thus, the brain treats two different types of negative events in distinct ways: glutamate inputs and GABA inputs may work together to produce inhibitory responses to reward omission, while GABA neurons may need to actively cancel out excitation by glutamate inputs to produce inhibitory responses to aversive stimuli in dopamine neurons (Figure 7G). These results are consistent with the idea that the inhibitory response to reward omission is a part of TD errors^4^, and thus already incorporated at the early stage of TD error computation, whereas the negative value of aversive events seems to be compiled into TD errors in later stages.

Competition between glutamate and GABA inputs in response to aversive stimuli raises the possibility that dopamine responses to aversive stimuli can be flexibly adjusted by modulating the balance of opposite information from excitatory and inhibitory inputs. One such example is opioid analgesia. While opioids affect multiple sites in pain pathways, as well as reward pathways, in the brain^63–65^, we found that buprenorphine, a partial μ-opioid agonist/κ-opioid antagonist, dramatically decreased inhibitory responses to an aversive stimulus in dopamine neurons but it did not affect their glutamate inputs. μ-opioid receptor expression is enriched in the RMTg and its direct and indirect upstream inputs such as the parabrachial nucleus (PBN), lHb, and the lateral preoptic area of the hypothalamus (LPO)^21,32–34,66^, and μ-opioid receptor activation inhibits signaling of these neurons^51–54^. Thus, it is highly probable that opioids preferentially decrease inhibitory input to dopamine neurons and tip the excitatory-inhibitory balance of inputs to dopamine neurons towards excitation (Figure 8F). We found that inhibitory responses of dopamine neurons to an aversive stimulus were greatly reduced or even flipped to excitation in some cases, consistent with the previous finding that ablation of RMTg flips dopamine responses to aversive electric shock from inhibition to excitation^26^. Importantly, dopamine inhibitory responses are thought to facilitate negative learning. Hence, opioids suppress dopamine signaling for negative learning with an aversive stimulus, which could contribute to analgesia as well as addiction by undermining the transmission of negative value information or negative RPEs of aversive events.

Our results demonstrate that glutamate and GABA neurons provide overlapping yet complementary information for dopamine neurons to compute TD error. Our quantification indicates that glutamate input contributes to a significant portion of dopamine TD errors, yet does not send negative value of aversive stimuli. These findings lend insight into the brain’s strategy for precise transmission of information of positive and negative valence: positive responses are generated by an increase of excitatory inputs, while negative responses are generated by an increase of inhibitory inputs^26,27,31^. A similar idea was proposed for pain responses where activation of ON cells or OFF cells, rather than activation and inhibition of ON cells, conveys pain information to inform behavioral decisions^67^. Following this framework, dopamine neurons may combine glutamate and GABA TD error information to overcome inherent constraints of each neurotransmitter (glutamate and GABA). For example, excitation of inputs, rather than inhibition, might be needed to produce moment-by-moment precise computation. If this is the case, it would be reasonable to use glutamate input preferentially for activation and GABA input preferentially for inhibition, especially by aversive stimuli, to produce intact TD errors in dopamine neurons^68^.

Excitatory-inhibitory parallel input systems might be functionally advantageous to generate dopamine activity. First, the combination of biased RPE inputs may provide flexibility in dopamine representation. For example, by changing the weights of glutamate and GABA inputs separately, dopamine neurons may provide positively or negatively biased teaching signals to other brain areas so that learning from positive and negative events is separately adjusted according to factors such as internal state and environmental context with other neuromodulators, hormone levels and neuropeptides such as endogenous opioids. Because both glutamate and GABA inputs send (biased) TD errors, these modulations may be able to shift dopamine responses positively or negatively, potentially even flipping responses to an aversive stimulus when appropriate in a given state without distorting core information (i.e. monotonic representation) of TD errors.

Second, different combinations of inputs may contribute to dopamine neuron diversity. For example, dopamine neurons that project to ventromedial VS are activated by aversive stimuli^69^. It has also been reported that dopamine axons in the dorsolateral striatum exhibit positively shifted RPE compared to dopamine axons in VS and dorsomedial striatum^46^. Because glutamate inputs convey positively shifted RPE and show excitation to aversive stimuli, differential weighting of inputs from glutamate neurons may partially explain reported diversity in dopamine activity. At a finer scale, a recent study proposed that the learning rate for negative and positive events is adjusted in each dopamine recipient neuron to represent the complete distribution of rewards, rather than simply expected value (i.e. distributional reinforcement learning^70,71^). Diverse weights of positively and negatively biased input onto dopamine neurons may provide raw material for learning such distributional representations.

Taken together, we found that TD error signals are already conveyed in the population activity of glutamate inputs, in contrast to existing models for TD error computation. Because fiber-fluorometry signals are the sum of signals across all inputs, the current observations cannot distinguish whether TD error computations took place at dopamine neurons or upstream. However, previous single neuron recording found that information for RPE in presynaptic neurons is mixed but not as complete as that conveyed by dopamine neurons, suggesting partial computation at multiple nodes, but completion at the level of dopamine neurons^22^. In the present study, we demonstrate that TD error signals carried by glutamate inputs are positively biased, and lack inhibitory responses to an aversive stimulus, strongly suggesting that mixing with GABA inputs is critical to form complete TD errors. While RPE is observed in various brain areas^22,25,28,45,72,73^, dopamine neurons may uniquely represent “complete” TD errors by mixing glutamate TD errors and GABA TD errors from distributed sources.

## Acknowledgments

We thank Shuhan Huang, Iku Tsutsui-Kimura, and Benedicte Babayan for technical assistance, Adam S. Lowet, Sara Matias, Isobel W. Green, Malcolm G. Campbell, and all lab members for discussion. We thank Catherine Dulac for sharing reagents and equipment. We thank Loren Looger, University of California, San Diego for pGP-AAV-CAG-FLEX-SF-iGluSnFR(A184S)-WPRE plasmid and AAV1-hSyn-SF-iGluSnFR(A184S), Douglas Kim and GENIE Project, Janelia Farm Research Campus, Howard Hughes Medical Institute for AAV8-hSyn-FLEX-jGCaMP7f, Edward Boyden, Media Lab, Massachusetts Institute of Technology for AAV5-CAG-FLEX-tdTomato and AAV8-hSyn-FLEX-ChrinsonR-tdTomato, and Yulong Li, State Key Laboratory of Membrane Biology, Peking University for AAV9-hSyn-DA2m. We thank the Harvard Center for Biological Imaging for infrastructure and support. This work was supported by grants from National Institute of Mental Health (R01MH125162, MW-U), National Institute of Health (U19 NS113201, NS 108740, NU), the Simons Collaboration on Global Brain (NU), Japan Society for the Promotion of Science, Japan Science and Technology Agency (RA).

## Author Contributions

R.A. and M.W.-U. designed experiments and analyzed data. R.A. collected data. The results were discussed and interpreted by R.A., N.U. and M.W.-U.. R.A. and M.W.-U. wrote the paper, and R.A., N.U. and M.W.-U. edited the paper.

## STAR Methods

### Animal

62 female and male mice, 2-19 months old were used in this study. We used heterozygote for DAT-Cre (Slc6a3^tm1.1(cre)Bkmn^; The Jackson Laboratory, 006660)^74^, LSL-tdTomato (Gt(ROSA)26Sor^tm14(CAG–tdTomato)Hze^; The Jackson Laboratory, 007914)^75^ transgenic lines, vGluT3-Cre (Slc17a8^tm1.1(cre)Hze^; The Jackson Laboratory, 028534)^76^ and homozygote for vGluT2-Cre (Slc17a6^tm2(cre)Lowl/J^; The Jackson Laboratory, 016963)^77^ line. DAT-Cre lines crossed with LSL-tdTomato were used for some sensor recordings. vGluT3-Cre lines crossed with LSL-tdTomato were used for some of the experiments with DA sensor, without use of Cre recombinase. Mice were housed on a 12 hr dark (7:00-19:00)/12 hr light (19:00-7:00) cycle. Experiments were performed in the dark period. Ambient temperature was kept at 23±2.7 C° and humidity was kept below 50%. All procedures were performed in accordance with the National Institutes of Health Guide for the Care and Use of Laboratory Animals and approved by the Harvard Animal Care and Use Committee.

### Virus

pGP-AAV-CAG-FLEX-SF-iGluSnFR(A184S)-WPRE (gift from Loren Looger; Addgene, #106186) was packaged into AAV at UNC vector core (AAV5-CAG-FLEX-SF-iGluSnFR(A184S)-WPRE, 2.4 × 10^13^ vg/ml; AAV8-CAG-FLEX-SF-iGluSnFR(A184S)-WPRE, 0.88 × 10^13^ and 1.7 × 10^13^ vg/ml). In addition to the above, the following AAVs were used in this study: AAV8-hSyn-FLEX-jGCaMP7f (gift from Douglas Kim & GENIE Project; 1.8 × 10^13^ vg/ml; Addgene, #104492-AAV8), AAV1-hSyn-SF-iGluSnFR(A184S) (gift from Loren Looger; 2.1× 10^13^ vg/ml; Addgene, #106174-AAV1), AAV9-hSyn-DA2m (gift from Yulong Li; 1.01 × 10^13^ vg/ml; ViGene bioscience), AAV5-CAG-FLEX-tdTomato (gift from Edward Boyden; 7.8 × 10^12^ vg/ml; UNC Vector Core), and AAV8-hSyn-FLEX-ChrinsonR-tdTomato (gift from Edward Boyden; 3.7 × 10^12^ vg/ml; UNC vector core).

### Surgery for virus injection, head-plate installation, and fiber implantation

The surgery was performed under aseptic conditions as previously described^30^. Mice were anesthetized with isoflurane (1-2% at 0.5-1 L/min) and local anesthetic (lidocaine (2%)/bupivacaine (0.5%) 1:1 mixture, S.C.) was applied at the incision site. Analgesia (ketoprofen for post-operative treatment, 5 mg/kg, I.P.; buprenorphine for pre-operative treatment, 0.1 mg/kg, I.P.) was administered for 3 days following surgery. A custom-made head-plate was placed on the well-cleaned and dried skull with adhesive cement (C&B Metabond, Parkell) containing a small amount of charcoal powder. For expression of glutamate sensor in the dopamine neurons, AAV8 (or AAV5)-CAG-FLEX-SF-iGluSnFR(A184S)-WPRE or mixture with AAV5-CAG-FLEX-tdTomato (2-8:1) was injected unilaterally in the VTA (600 nl, Bregma -3.05-3.15 mm AP, 0.375-0.75 mm ML, 4.35 mm DV from dura) in DAT-Cre/LSL-tdTomato mice or DAT-Cre mice, respectively. For expression of GCaMP in dopamine neurons, AAV8-hSyn-FLEX-jGCaMP7f or mixed solution with AAV5-CAG-FLEX-tdTomato (2:1) was injected unilaterally in the VTA (500 nl, Bregma -3.05 mm AP, 0.325-0.75 mm ML, 4.35 mm DV from dura) in DAT-Cre/LSL-tdTomato mice or DAT-Cre mice, respectively. For expression of dopamine sensor in the VS, AAV9-hSyn-DA2m was injected unilaterally in the VS (300 nl, Bregma +1.45 AP, 1.4 ML, 4.35 DV from dura) in DAT-Cre/tdTomato mice or vGluT3-Cre/LDL-tdTomato mice. For optogenetic stimulation experiment, AAV8-hSyn-FLEX-ChrinsonR-tdTomato was injected unilaterally in the VP, STh, and PPTg (VP: 300 nl, 20° angled (tip is directed to medial), Bregma +1.0 AP, 2.5 ML, 4.2 DV from dura; STh: 300 nl, Bregma - 2.2 AP, 1.5 ML, 4.5 DV from dura; PPTg: 300 nl, 25° angled (tip is directed to medial), Bregma -4.75 AP, 2.35 ML, 4.5 DV from dura) in vGluT2-Cre mice to express ChrimsonR in glutamate input to the VTA, and AAV1-hSyn-SF-iGluSnFR(A184S)/AAV5-CAG-FLEX-tdTomato (8:3) or AAV9-hSyn-DA2m was injected in the VTA (250 nl, Bregma -3.1 mm AP, 0.5 mm ML, 4.35 mm DV from dura) or VS (300 nl, Bregma +1.45 AP, 1.4 ML, 4.2 DV from dura), respectively. A glass pipette containing AAV was slowly moved down to the target over the course of a few minutes and kept for 2 minutes to make it stable. AAV solution was slowly injected (∼15 min) and the pipette was left for 10-20 min. Then the pipette was slowly removed over the course of several minutes to prevent the leak of virus and damage to the tissue. An optical fiber (400 μm core diameter, 0.66 or 0.48 NA; Doric) was implanted in the VS (Bregma +1.45 mm AP, 1.4 mm ML, 4.1-4.0 mm DV from dura) or the VTA (Bregma -3.05 mm AP, 0.325-0.75 mm ML, 4.15-4.3 mm DV from dura). For the optogenetic stimulation, stimulation fiber was implanted in the VTA (15° angled (tip is directed to caudal), Bregma -1.85 mm AP, 0. 5 mm ML, 4.1 mm DV from dura), and recording fiber was implanted in the VTA (Bregma -3.1 mm AP, 0.5 mm ML, 4.2 mm DV from dura) or VS (15° angled (tip is directed to caudal), Bregma +2.7 AP, 1.4 ML, 4.35 DV from dura), respectively. The fiber was slowly lowered to the target and fixed with adhesive cement (C&B Metabond, Parkell) containing charcoal powder to prevent contamination of environmental light and leak of laser light. A small amount of rapid-curing epoxy (Devcon, A00254) was applied on the cement to glue the fiber better.

### Fiber-fluorometry (photometry)

To effectively collect the fluorescence signal from the deep brain structure, we used custom-made fiber-fluorometry as previously described^30,41^. Blue light from 473 nm DPSS laser (Opto Engine LLC) and green light from 561 nm DPSS laser (Opto Engine LLC) were attenuated through neutral density filter (4.0 optical density, Thorlabs) and coupled into an optical fiber patch cord (400 μm, Doric) using 0.65 NA 20x objective lens (Olympus). This patch cord was connected to the implanted fiber to deliver excitation light to the brain and collect the fluorescence emission signals from the brain simultaneously. The green and red fluorescence signals from the brain were spectrally separated from the excitation lights using a dichroic mirror (FF01-493/574-Di01, Semrock). The fluorescence signals were separated into green and red signals using another dichroic mirror (T556lpxr,Chroma), passed through a band pass filter (ET500/40x for green, Chroma; FF01-661/20 for red, Semrock), focused onto a photodetector (FDS10X10, Thorlabs), and connected to a current amplifier (SR570, Stanford Research systems). The preamplifier outputs (voltage signals) were digitized through a NIDAQ board (PCI-e6321, National Instruments) and stored in a computer using custom software written in LabVIEW (National Instruments). The sampling rate was 1000 Hz. Light intensity at the tip of patch cord was adjusted to 100-200 μW (glutamate sensor and GCaMP) and 50 μW (dopamine sensor).

### Optogenetic stimulation

Red light from 625 nm LED light (M625F2, Thorlab) was applied through an optical patch cord (400 μm, 0.39 NA, Thorlab). 12 mW single block pulse light of 250ms duration was triggered through custom software written in LabVIEW (National Instruments) via a NIDAQ board (PCI-e6321, National Instruments). A variable inter-trial interval (ITI) of flat hazard function (minimum 10s, mean 13s, truncated at 20s) was placed between trials.

### Histology

All mice used in the experiments were examined for histology to confirm the fiber position as previously described^30^. The mice were deeply anesthetized by an overdose of ketamine/medetomidine, exsanguinated with phosphate buffered saline (PBS), and perfused with 4% paraformaldehyde (PFA) in PBS. The brain was dissected from the skull and immersed in the 4% PFA for 12-24 hours at 4 °C. The brain was rinsed with PBS and sectioned (100 μm) by vibrating microtome (VT1000S, Leica). Immunohistochemistry with TH antibody (AB152, Millipore Sigma; 1/750) was performed to identify dopamine neurons, with GFP antibody (GFP-1010, Aves Labs; 1/3000) to localize sensor-expressing areas when GCaMP and glutamate sensor raw signals were not strong enough. The sections were mounted on a slide-glass with a mounting medium containing 4’,6-diamidino-2-phenylindole (VECTASHIELD, Vector laboratories) and imaged with Axio Scan.Z1 (Zeiss) or LSM880 with FLIM (Zeiss).

### Behavior

After 1 week of recovery from surgery, mice were water-restricted in their cages. All conditioning tasks were controlled by a NIDAQ board and LabVIEW. Mice were handled for 2 days, acclimated to the experimental setup for 1-2 days including consumption of water from the tube, and head-fixed with random interval water for 1-3 days until mice show reliable water consumption. For odor-based classical conditioning, all mice were head-fixed, and the volume of water reward was constant for all reward trials (predicted or unpredicted) in all conditions (6 μl). Some sessions included mild air puff trials, directed at one of the eyes and the intensity of air puff was constant for all air puff trials (predicted or unpredicted; 2.5 psi). In classical conditioning, each association trial began with an odor cue (lasting 1 s) followed by a 2 s delay, and then an outcome (either water, nothing, or air puff) was delivered. In sequential conditioning, each association trial began with a distal odor cue (lasting 1 s) followed by a 2 s delay, and then a distal odor cue (lasting 1 s) followed by 1 s delay, and then an outcome (either water, or nothing) was delivered. Some trial types began with a proximal odor cue (lasting1 s) followed by 1 s delay, and then an outcome (either water, or nothing) was delivered. Odors were delivered using a custom olfactometer.^78^ Each odor was dissolved in mineral oil at 1:10 dilution and 30 μl of diluted odor solution was applied to the syringe filter (2.7μm pore, 13mm; Whatman, 6823-1327). Odorized air was further diluted with filtered air by 1:8 to produce a 900 ml/min total flow rate. Different sets of odors (Ethyl butyrate, p-Cymene, Isoamyl acetate, Isobutyl propionate, 1-Butanol, 4-Methylanisole, Caproic acid, Eugenol, and 1-Hexanol) were selected for each animal. Some of the animals shared the same odor set (4 animals and 3 animals for DA sensor classical conditioning; 3 animals and 2 animals for sequential conditioning)

A variable inter-trial interval (ITI) of flat hazard function (minimum 10s, mean 13s, truncated at 20s) was placed between trials. Each session was composed of multiple blocks (12-24 trial/block) and all trial types were pseudorandomized in each block. Each day, the mice did about 70-350 trials over the course of 20-75 min, and with constant excitation from the laser and continuous recording in recording sessions.

Training for classical conditioning (Figures 2, 4 and 6) used 4 types of trials; odor cue predicting 100% water, odor cue predicting 40% water/60% no outcome (nothing), odor cue predicting nothing (29.4% of all trials for each odor), and water without cue (free water) (11.8%) for 7-10 days, and then odor cue predicting 80% water/20% nothing, odor cue predicting 40% water/60% nothing, odor cue predicting nothing (29.4% each), and free water (11.8%) for more than 2 days of training followed by recording sessions. 3 animals used for glutamate sensor recording (Figure 2) were trained with classical conditioning with air puff trial; odor cue predicting 100% water, odor cue predicting 100% air puff, odor cue predicting 40% water/60% no outcome (nothing), odor cue predicting nothing (20.8 % of all trials for each odor), and water without cue (free water), air puff without cue (free air puff) (8.3 % of all trials for each stimulus) for 8-9 days, and then odor cue predicting 80% water/20% nothing, odor cue predicting 80% air puff/20% nothing, odor cue predicting 40% water/60% nothing, odor cue predicting nothing (20.8 % each), and water without cue (free water), air puff without cue (free air puff) (8.3% each) for more than 7 days of training followed by recording sessions. Of note these 3 animals are used only in the analysis for glutamate sensor activity pattern (Figure 2), and not included for comparison between different sensor recordings (Figures 4 and 6).

For sequential conditioning step1 (Figure 3), mice were trained with proximal odor for 3-7 days, using 3 types of trials; proximal odor cue predicting 100% water, proximal odor cue predicting nothing (45.8 % of all trials for each odor), and water without cue (free water) (8.3 %). Then, for sequential conditioning step2, mice were further trained with distal odor and proximal odor using 6 types of trials; distal odor cue predicting proximal odor cue predicting 100% water (25 %), distal odor cue predicting 50% proximal odor cue predicting water/50% proximal odor cue predicting nothing (25 %), distal odor cue predicting proximal odor cue predicting nothing (25 %), proximal odor cue predicting 100% water (8.3 %), proximal odor cue predicting nothing (8.3 %), and water without cue (free water) (8.3 %). Sensor signals were recorded after 8-12 days of step2 training.

For buprenorphine treatment, the mouse received unexpected air puff and water with a variable ITI of the flat hazard function (minimum 10s, mean 13s, truncated at 20s). Animals were first acclimated to head-fixation with water reward for 2 days. Then the animals were habituated to a test procedure for two days, which was composed of 40-70 trials of air puff and water presented in pseudorandomized order (the order of trials is pseudorandomized within each block of 10 trials composed of the same number of air puff trials and reward trials) with subcutaneous injection of saline (10 μl/g in body weights) at the end of sessions. After habituation sessions, buprenorphine sessions and control saline sessions were performed in pseudorandom order. Each session was separated into pre-injection sub-session and post-injection sub-session. The sub-session was composed of 48-75 trials with the air puff and water trials in the pseudorandom order. After pre-injection sub-sessions, either buprenorphine (Buprenorphine hydrochloride, Par Pharmaceutical: diluted in saline 0.03 mg/ml) or control saline was subcutaneously injected (10 μl/g; 0.3 μg/g buprenorphine or corresponding volume of saline). Post-injection sub-session started 5 min after injection. To prevent acute adverse effects, we used a smaller dose of buprenorphine (0.15 μg/g) in the first session and this session is not included in the analysis.

### Data analysis

#### Fiber-fluorometry

The noise from the power line in the voltage signal was cleaned by removing 58-62Hz signals through a band stop filter. The signal was smoothed with moving average of 50 ms and the global change within a session was normalized using a moving median of 100 s. Then, the correlation between green and red signals during ITI was examined by linear regression. If the correlation is significant (p<0.05), fitted tdTomato signals were subtracted from green signals. Responses were calculated by subtracting the average baseline activity from the average activity of the target window. Z-scores of the signals were obtained using mean and standard deviation of signals in all trials (from 1 s before odor onset to 8 s after odor onset for classical conditioning, from 1 s before distal odor onset to 13 s after odor onset for sequential conditioning) in each animal. Stimulus responses were measured as average activity of analysis window (1-1000 ms for odor response in classical conditioning, 1-1000 ms for distal odor response in sequential conditioning, 201-1200 ms for proximal odor response in sequential conditioning, 201-1200 ms for water and air puff response, 1501-2500 ms for omission response, 1-500 ms for optogenetic stimulation response).

#### Quantification of rise and decay of sensor signals

Rise time of glutamate sensor and dopamine sensor signal pattern for free water and optogenetic stimulation was defined as latency from 10% peak signal timing to 90% peak signal timing. Decay time was defined as latency from peak timing to 50% peak activity timing. The activity peak during response period (3 sec from the stimulus onset) was detected by finding a maximum response in moving windows of 20 ms that exceeds 2 × standard deviation of baseline activity (moving windows of 20 ms during -1 to 0 sec from an odor onset).

#### Test for monotonic relationship between neural responses and reward outcome

The responses in the sensor signals were normalized by the average responses to chosen events in one session as following: 80% reward odor for odor responses in classical conditioning (1-1000 ms), odor 1 for distal odor in sequential conditioning (1-1000 ms), and unexpected odor A for proximal odor in sequential conditioning (201-1200 ms). Responses in each trial were fitted with reward outcome with linear regression for each animal. Zero-crossing points were determined as x-intercepts of obtained linear line to estimate reward probability that produce zero response in sensor signals by odor in classical conditioning and distal odor in sequential conditioning for each animal.

#### Estimation of dopamine sensor signal based on the glutamate sensor signal

To estimate dopamine sensor signals from the glutamate sensor signals, we deconvolved glutamate sensor signals in sequential conditioning in each animal (average in 3 sessions) with “glutamate kernels” (interval of 200 ms) using optogenetic stimulation responses in glutamate sensor signals (0-3.5 s)^79^, and then convolved the resulting trace with “dopamine kernels” using optogenetic responses in dopamine sensor signals (0-3.5 s) (Figure 5C). We used down-sampled (every 20 ms) responses in all trials for the model fitting. We used different kernels for each trial type (Figure 3A). Odor kernels consist of 6 types of kernels: ‘Odor1-OdorA-water’, ‘Odor2-OdorA-water’, ‘Odor2-OdorB-nothing’, and ‘Odor3-OdorB-nothing’ kernels to span 0 to 5 s from distal odor onset, and ‘No odor-OdorA-water’ and ‘No odor-OdorB-nothing’ kernels to span 0 to 2 s from proximal odor onset. Water kernels consist of 2 types of kernels: ‘expected water’ kernels in trials with water after odorA, and ‘free water’ kernels in trials with unexpected water to span 0–4 s from water onset. Glutamate sensor responses were fitted with all the kernels using linear regression with Elastic net regularization (α=0.75) with 10-fold cross validation. The regularization coefficient lambda was chosen so that cross-validation error is minimum plus one standard deviation. Glutamate kernels were swapped with “dopamine kernels” using optogenetic responses in dopamine sensor signals. The amplitude of the obtained trace was adjusted by linear regression with DA sensor responses in free water trials. % explained by an estimated DA senor signal was expressed as reduction of a variance in the residual responses compared to the recorded DA sensor responses.

#### Quantification of the effect of buprenorphine administration

Air puff response (1-1000ms) was quantified as the average of 4 sessions for each animal for each condition in 6 animals. We used fewer sessions for two animals because of weak signals (peak of water response below 2 z-score), or system problems (1 buprenorphine session and 2 saline sessions for one animal and 3 buprenorphine sessions and 4 saline sessions for one animal). We used all trials of all sessions to test the significance of the response for each animal (Figures 8B and 8D).

#### Licking

Licking from a water spout was detected by a photoelectric sensor that produces a change in voltage when the light path is broken. The timing of each lick was detected at the peak of the voltage signal above a threshold. To plot the time course of licking patterns, the lick rate was calculated by a moving average of a 200 ms window.

### Statistics & Reproducibility

All analyses were performed using custom software written in Matlab (MathWorks). All statistical tests were two-sided. A boxplot indicates 25th and 75th percentiles as the bottom and top edges, respectively. The center line indicates the median. The whiskers extend to the most extreme data that is not considered as outlier. In other graphs, error bars show standard errors. To test for significance of the model fitting, the *p*-value for the F-test on the model was calculated.

One-sample *t*-test was performed to test if the mean of a data set is not equal to zero. To compare the difference of the mean between two groups, two-sample *t*-test was performed. To compare the difference of the mean between pairs of measurement, paired *t*-test was performed. *p*-value less than or equal to 0.05 was regarded as significant for all tests. Asterisks in the figures stand for *** *p* ≤ 0.001, ** *p* ≤ 0.01, and * *p* ≤ 0.05. “n.s.” stands for *p* > 0.05. For comparison of more than three measurements, one-way ANOVA followed by Tukey’s test was performed, and when the measurement contains two independent variables, two-way ANOVA was performed. The two-sample Kolmogorov-Smirnov test was used to compare distribution of response changes (Figure 8E). Data distribution was assumed to be normal, but this was not formally tested. No formal statistical analysis was carried out to predetermine sample sizes but our sample sizes are similar to those reported in previous publications^41,46,80^. No animals were excluded from the study: all analysis includes data from all animals.

### Lead contact

Further information and requests for resources and reagents should be directed to and will be fulfilled by the lead contact, Mitsuko Watabe-Uchida.

### Materials availability

This study did not generate new unique reagents.

### Data availability

The fluorometry data will be shared at a public deposit.

### Code availability

All conventional codes used to obtain the result will be available from a public deposit source.

---

**Supplemental Figure 1.**
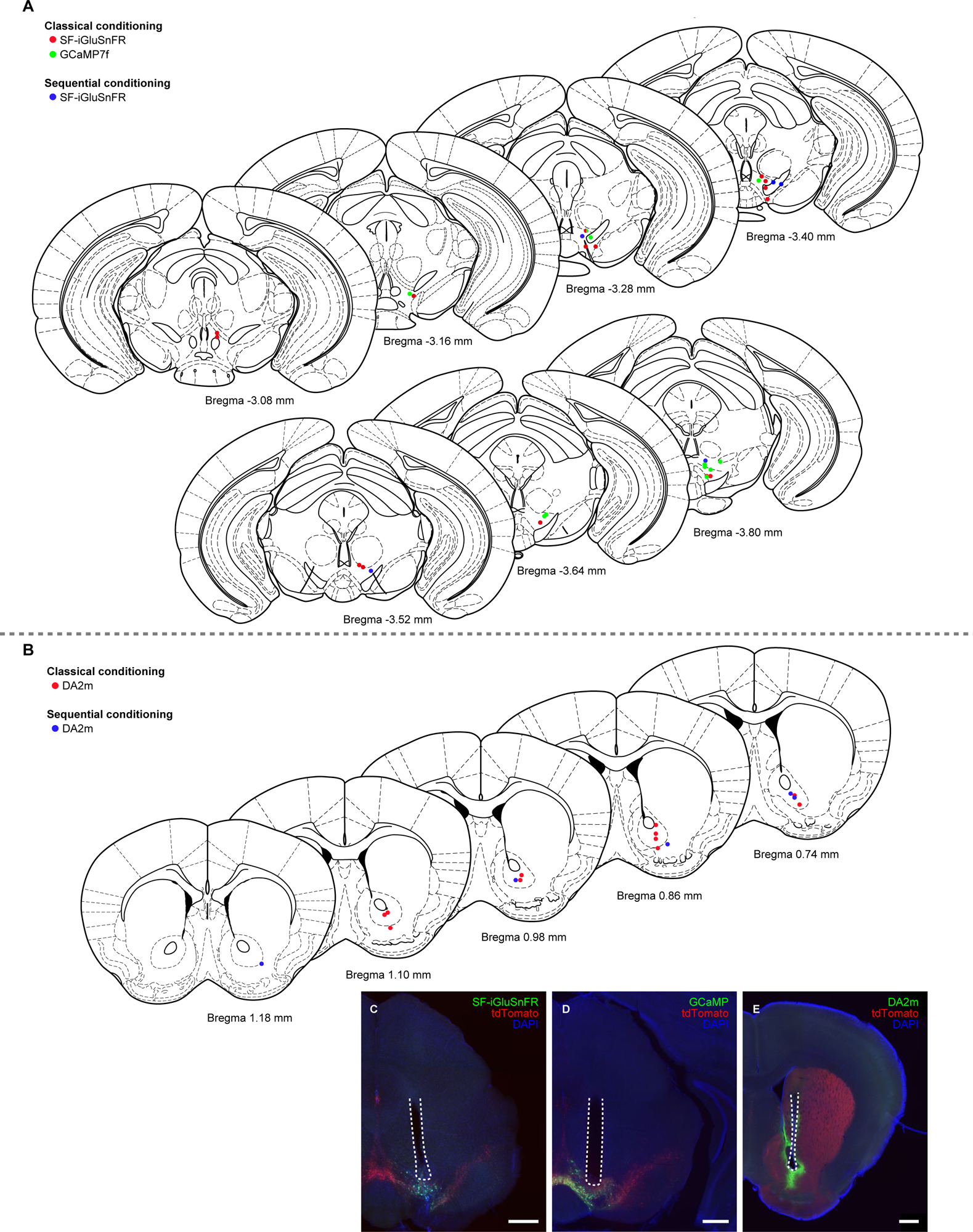
Recording sites in classical conditioning and sequential conditioning. (A) Recording sites in the VTA for glutamate sensor (SF-iGluSnFR: red for classical conditioning, 15 animals recorded from left hemisphere; blue for sequential conditioning, 5 animals recorded from left hemisphere) and calcium sensor (GCaMP7f: green for classical conditioning, 11 animals recorded from left hemisphere) expressed in VTA dopamine neurons. (B) Recording sites in the VS for dopamine sensor (DA2m: red for classical conditioning, 7 animals recorded from left hemisphere and 4 animals recorded from right hemisphere; blue for sequential conditioning, 5 animals recorded from the right hemisphere) expressed in striatal cells. (C-E) Example recording sites for glutamate sensor (C, green), calcium sensor (D, green), and dopamine sensor (E, green) recording site. The dotted lines are optic fiber tracks. tdTomato (red) is expressed under the control of DAT-Cre. Counterstained with DAPI (blue). Scale bars: 500 μm.

**Supplemental Figure 2.**
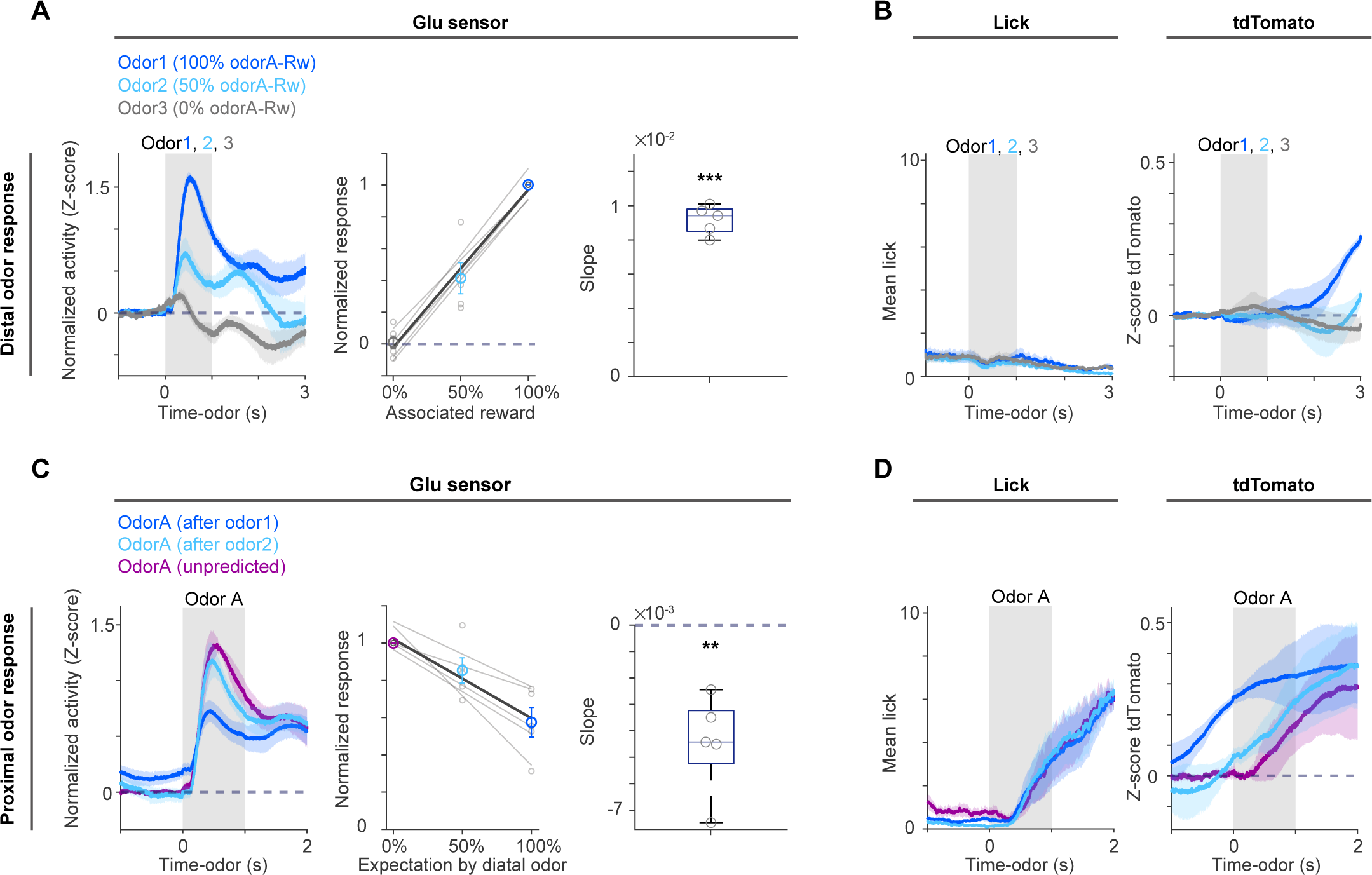
Glutamate sensor signal, control tdTomato signal, and licking during sequential conditioning. (A) Glutamate sensor signal at distal odor cue without noise-correction (n = 5 animals). Left, mean ± s.e.m. Middle and right, linear regression of responses to distal odors (1-1000 ms from distal odor onset) with associated reward probability. Middle, light gray, each animal; dark color, average of all animals, correlation coefficient 0.95, *p* = 0.42×10^-7^, *F*-test; mean ± s.e.m.. Right, regression coefficients in each animal (*t* = 24, *p* = 0.16×10^-4^, two-sided *t*-test). (B) Lick (left) and control tdTomato signal (right) at distal odor cue. Mean ± s.e.m. (C) Glutamate sensor signal at odor A (n = 5 animals) without noise-correction. Left, mean ± s.e.m. Middle and right, linear regression of responses to odor A (201-1200 ms from odor A onset) with expectation of odor A. Middle, light gray, each animal; dark color, average of all animals, correlation coefficient -0.81, *p* = 0.23×10^-3^, *F*-test; mean ± s.e.m.. Right, regression coefficients in each animal (*t* = -5.3, *p* = 0.59×10^-2^, two-sided *t*-test). (D) Lick (left) and control tdTomato signal (right) at odor A. Mean ± s.e.m. In box plots, grey lines are the median; edges are 25^th^ and 75^th^ percentiles; and whiskers are the most extreme data points not considered as outliers.

**Supplemental Figure 3.**
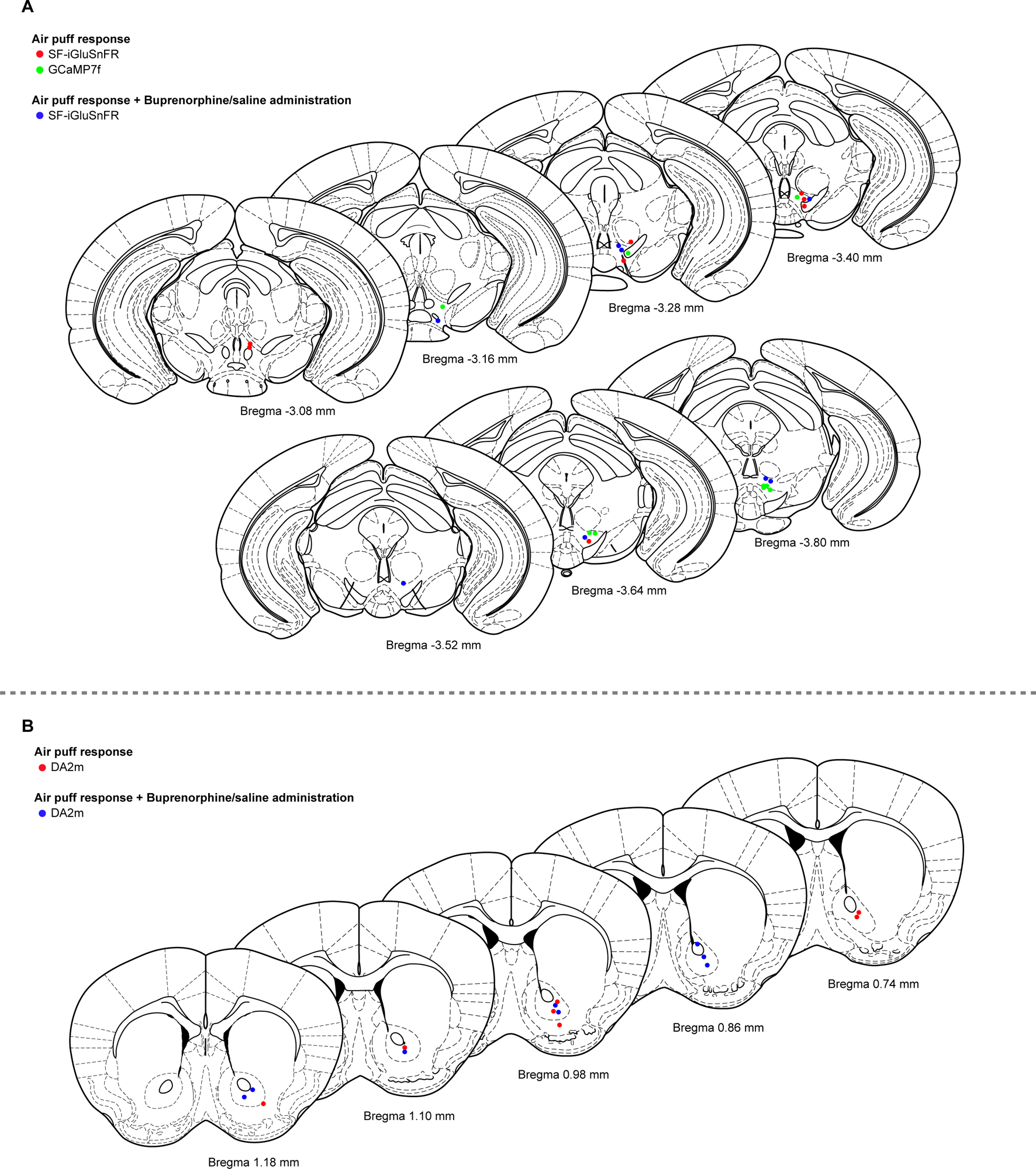
Recording sites for air puff response with or without buprenorphine treatment. (A) Recording sites in the VTA for glutamate sensor (SF-iGluSnFR: red for no treatment, 8 animals recorded from left hemisphere; blue for buprenorphine treatment, 8 animals recorded from left hemisphere) and calcium sensor (GCaMP7f: green for no treatment, 9 animals recorded from left hemisphere) expressed in VTA dopamine neurons. (B) Recording sites in the VS for dopamine sensor (DA2m: red for no treatment, 7 animals recorded from right hemisphere; blue for buprenorphine treatment, 8 animals recorded from right hemisphere) expressed in striatal cells.

